# DNA Methylation Atlas of the Mouse Brain at Single-Cell Resolution

**DOI:** 10.1101/2020.04.30.069377

**Authors:** Hanqing Liu, Jingtian Zhou, Wei Tian, Chongyuan Luo, Anna Bartlett, Andrew Aldridge, Jacinta Lucero, Julia K. Osteen, Joseph R. Nery, Huaming Chen, Angeline Rivkin, Rosa G Castanon, Ben Clock, Yang Eric Li, Xiaomeng Hou, Olivier B. Poirion, Sebastian Preissl, Carolyn O’Connor, Lara Boggeman, Conor Fitzpatrick, Michael Nunn, Eran A. Mukamel, Zhuzhu Zhang, Edward M. Callaway, Bing Ren, Jesse R. Dixon, M. Margarita Behrens, Joseph R. Ecker

## Abstract

Mammalian brain cells are remarkably diverse in gene expression, anatomy, and function, yet the regulatory DNA landscape underlying this extensive heterogeneity is poorly understood. We carried out a comprehensive assessment of the epigenomes of mouse brain cell types by applying single nucleus DNA methylation sequencing to profile 110,294 nuclei from 45 regions of the mouse cortex, hippocampus, striatum, pallidum, and olfactory areas. We identified 161 cell clusters with distinct spatial locations and projection targets. We constructed taxonomies of these epigenetic types, annotated with signature genes, regulatory elements, and transcription factors. These features indicate the potential regulatory landscape supporting the assignment of putative cell types, and reveal repetitive usage of regulators in excitatory and inhibitory cells for determining subtypes. The DNA methylation landscape of excitatory neurons in the cortex and hippocampus varied continuously along spatial gradients. Using this deep dataset, an artificial neural network model was constructed that precisely predicts single neuron cell-type identity and brain area spatial location. Integration of high-resolution DNA methylomes with single-nucleus chromatin accessibility data allowed prediction of high-confidence enhancer-gene interactions for all identified cell types, which were subsequently validated by cell-type-specific chromatin conformation capture experiments. By combining multi-omic datasets (DNA methylation, chromatin contacts, and open chromatin) from single nuclei and annotating the regulatory genome of hundreds of cell types in the mouse brain, our DNA methylation atlas establishes the epigenetic basis for neuronal diversity and spatial organization throughout the mouse brain.

## Introduction

Epigenomic dynamics is associated with cell differentiation and maturation in the mammalian brain, and plays an essential role in regulating neuronal functions and animal behavior^1-3^. Cytosine DNA methylation (5mC) is a stable covalent modification that persists in post-mitotic cells throughout their lifetime, and is critical for proper gene regulation^2,4,5^. In mammalian genomes, cytosine methylation predominantly occurs at CpG sites, showing dynamic patterns at regulatory elements with tissue and cell-type specificity^2,6-8^, modulating binding affinity of transcription factors (TF)^9,10^, and controlling gene transcription^11^. As a unique signature of neuronal cells, non-CpG (CH) cytosines are also abundantly methylated in the mouse and human brain^2^, which can directly affect DNA binding of MeCP2 methyl-binding protein responsible for Rett Syndrome^12-14^. mCH levels at gene bodies are anti-correlated with gene expression and show a remarkable diversity across neuronal cell types^7,8^.

Deeper understanding of epigenomic diversity in the mouse brain not only provides a complementary approach to transcriptome-based profiling methods to identify all brain cell types but additionally allows genome-wide prediction of the regulatory DNA sequences and transcriptional networks that underlie this diversity. Our previous studies demonstrated the utility of studying brain cell-type and regulatory diversity using single nucleus methylome sequencing (snmC-seq)^8,15^. In this study, we use snmC-seq2^16^ to conduct thorough methylome profiling with detailed spatial dissection in the adult postnatal day 56 (P56) mouse brain (Fig. 1a). In Li et al. (Companion Manuscript # 11), the same tissue samples were also profiled in parallel using single nucleus ATAC-seq (snATAC-seq) to identify genome-wide accessible chromatin^17^, providing complementary epigenomic information to aid in cell-type specific regulatory genome annotation. Moreover, to further study cis-regulatory elements and their potential targeting genes across the whole genome, we applied a multi-omic assay sn-m3C-seq^18^ to profile methylome and chromatin conformation in the same cells.

**Figure 1.**
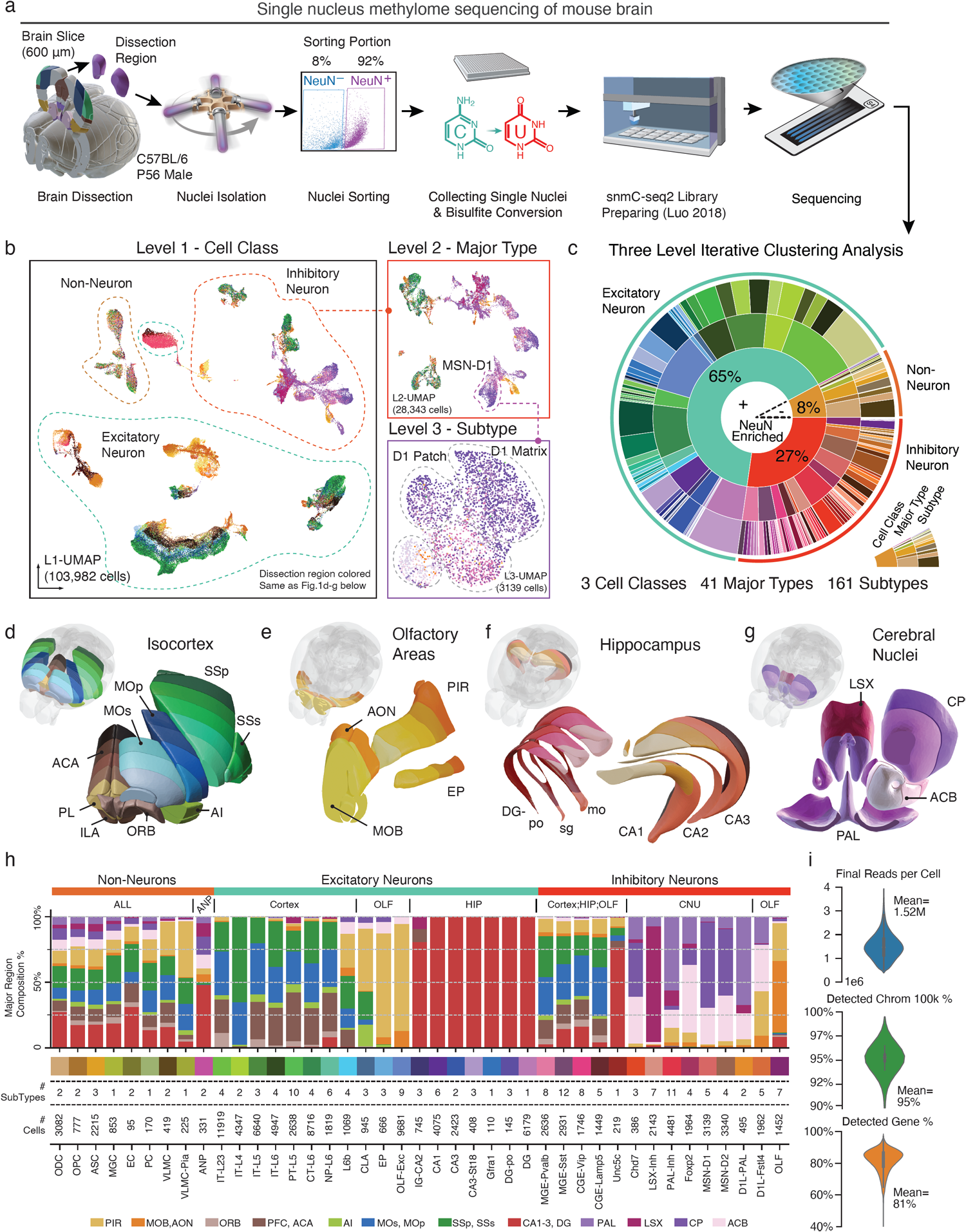
A survey of single-cell DNA methylomes in the mouse brain. **a**, The workflow of dissection, FANS, and snmC-seq2 library preparation. **b**, Level1 UMAP (L1-UMAP) is colored by the dissection region; the color palette is the same as (**d-g**). Panels show examples of Level 2 and Level 3 UMAP embeddings from iterative analyses. **c**, Sunburst of three-level iterative clustering. From inner to outer: L1 Cell Classes, L2 Major Types, and L3 Subtypes. **d-g**, Brain 3D dissection models of Isocortex (**d**), olfactory areas (**e**), hippocampus (**f**), and cerebral nuclei (**g**) colored by 45 dissection regions (see Extended Data Fig. 1 for the detail labels). **h**, An integrated overview of brain region composition, subtype, and cell numbers of the major types. **i**, The number of final pass QC reads, the percentage of nonoverlapping chromosome 100kb bins detected, and the percentage of GENCODE vm22 genes detected per cell.

The deep epigenomic datasets described here provide a detailed and comprehensive census of the diversity of cell-types across mouse brain regions, allowing prediction of cell-type-specific regulatory elements as well as their candidate target genes and upstream TFs. Highlights of the findings of our study include:

- Single nucleus methylome sequencing of 110,294 cells from the 45 dissected regions across cortex, hippocampus, striatum, pallidum, and olfactory areas of the mouse brain provides the largest single-cell base-resolution DNA methylation dataset.
- A robust single cell methylome analysis framework identifies 161 predicted mouse brain subtypes. The relationship between subtypes in distinct brain areas provides novel insights into anatomical cell/sub-structure associations.
- Comparing subtype-level methylomes allows identification of 3.5M cell-type-specific differentially CG methylated regions (CG-DMR) that cover ∼ 50% (1,240 Mb) of the mouse genome.
- Differentially methylated TF genes and binding motifs can be associated with branches in subtype phylogeny, allowing the prediction of cell-type gene regulatory programs specific for each developmental lineage.
- Integrative analysis with chromatin accessibility based cell clusters validate most methylome-derived subtypes, allowing prediction of 1.6M enhancer-like DMRs (eDMR) across cell subtypes.
- Identify cis-regulatory interactions between enhancers and genes with computational prediction and single-cell chromatin conformation profiling (in hippocampus).
- A continuous three-dimensional spatial methylation gradient was observed in cortical excitatory and dentate gyrus granule cells, with region-specific TFs and motifs found associated with the gradients.
- An artificial neural network model constructed using this deep dataset precisely predicts single neuron cell-type identity and brain area spatial location using only its methylome profile as input.

## Results

### A single cell DNA methylome atlas of 45 mouse brain regions

We used single-nucleus methylcytosine sequencing 2 (snmC-seq2)^16^ to profile genome-wide cytosine DNA methylation at single-cell resolution across four major brain structures: cortex, hippocampus (HIP), striatum and pallidum (or cerebral nuclei, CNU), and olfactory areas (OLF) (Fig. 1d-g) using adult male C57BL/6 mice (age P56-63). In total we analyzed 45 dissected regions in two replicates (Extended Data Fig. 1, Supplementary Table 2). Fluorescence-activated nuclei sorting (FANS) of antibody-labeled nuclei was applied to capture NeuN positive (NeuN^+^, 92%) neurons, while also sample a smaller number of NeuN negative (NeuN^-^, 8%) non-neuronal cells (Fig. 1a). In total, we profiled the DNA methylomes of 110,294 single nuclei yielding, on average, 1.5 million stringently filtered reads/cell (mean ± SD: 1.5×10^6^ ± 5.8×10^5^), covering 6.2 ± 2.6% of the mouse genome per cell. In each cell, we reliably quantified DNA methylation for 95 ± 4% of the mouse genome in 100kb bins and 81 ± 8% of gene bodies (Fig. 1i). In parallel, we performed single-nucleus ATAC-seq (snATAC-seq)^17^ on nuclei from the same samples to identify sites of accessible chromatin (see below and Li et al., companion manuscript # 11).

Based on the DNA methylome profiles in both CpG sites (CG methylation or mCG hereafter) and non-CpG sites (CH methylation or mCH) in 100kb bins throughout the genome, we performed a three-level iterative clustering analysis to categorize the epigenomic cell populations (Fig. 1b, c). After quality control and preprocessing (see Methods), in the first level (cell class), we clustered 103,982 cells into 67,472 (65%) excitatory neurons, 28,343 (27%) inhibitory neurons, and 8,167 (8%) non-neurons. The portion of non-neurons is determined by the FANS of NeuN^-^ nuclei (Supplementary Table 3, Extended Data Fig. 2b). The second round of iterative analysis of each cell class identified 41 major types in total (cluster size range 95-11,919, median 1,819 cells), and the third round separated these major types further into 161 cell subtypes (cluster size range 12-6,551, median 298 cells). Each subtype can be distinguished from all others based on: 1) a supervised model that reproduces the cluster label with > 0.9 cross-validated accuracy; and 2) at least 5 differentially methylated genes between each pair of subtypes (see Methods, and Extended Data Fig. 2, Supplementary Table 4 for subtype names and signature genes). All subtypes are highly conserved across replicates (Extended Data Fig. 2d), and replicates from the same brain region are co-clustered compared to samples from other brain regions (Extended Data Fig. 2e-g).

The spatial distribution of each cell type is assessed based on where the cells were dissected from (Supplementary Table 6). Here we used the Uniform Manifold Approximation and Projection (UMAP)^19^ to visualize epigenetic differences based on cell location (Fig. 1b, Extended Data Fig. 3) and major cell type (Extended Data Fig. 2a). Major non-neuronal types have a similar distribution across brain regions (Fig. 1h), with the exception of Adult Neuron Progenitors (ANP). We found two subtypes of ANP presumably corresponding to neuron precursors in the subgranular zone of the dentate gyrus (DG)^20^ (“ANP anp-dg”, 121 cells) and the rostral migratory stream (RMS)^21^ in CNU and OLF (“ANP anp-olf-cnu”, 210 cells). Major excitatory neuron types from isocortex, OLF, and HIP formed separate subtypes, with some exceptions potentially due to overlaps in dissected regions (see “Potential Overlap” column in Supplementary Table 2). Cells from isocortex were assigned to subtypes based on their projection types^8,22,23^. The Intratelencephalic (IT) neurons from all cortical regions contain four major types corresponding to the laminar layers (L2/3, L4, L5, and L6), with additional subtypes in each layer that differ across cortical regions. We also identified major types from cortical subplate structures, including the claustrum (CLA) and endopiriform nucleus (EP) from isocortex and OLF dissections.

GABAergic inhibitory neurons from isocortex and HIP cluster together into five major types, but most HIP neurons are then separated from isocortical neurons at the subtype level, in agreement with a companion transcriptomic study^24^. Intriguingly, in one major GABAergic type marked by low methylation at the gene *Unc5c* (suggesting high RNA expression), cells from HIP and isocortex remain to be co-clustered at subtype levels. The isocortex cells in this cluster display hypo-mCH in the gene body of both *Lhx6* and *Adarb2*, corresponding to the previously reported “Lamp5-Lhx6” cluster in transcriptome studies^22,25,26^. The close relationship between this cell type and HIP interneurons was also reported in a single-cell transcriptomic study in marmoset^27^. In contrast with excitatory neurons, subtypes of isocortical GABAergic neurons do not separate by cortical region. Interneurons from CNU and OLF group into nine major types and many subtypes which further separate by region (see vignette below), indicating substantial spatial diversity among CNU and OLF interneurons. In the next section, we provide several analytic vignettes to illustrate the unprecedented level of neuronal subtype and spatial diversity observed in their DNA methylomes.

### Neuron subtype vignettes

#### Methylome similarity between lndusium Griseum and Hippocampal region CA2 neuron subtypes

Nearly all hippocampal excitatory neurons formed their own clusters separately from cells in other brain regions (Fig. 1h), except the IG-CA2 neurons (745 cells, three subtypes; IG: Indusium griseum, CA: Cornu Ammonis) (Fig. 2c). Several markers of the hippocampal region CA2^28^ (e.g., *Pcp4*^*29*^, *Cacng5*^*30*^, *Ntf3*^*31*^) were marked by low gene body mCH (Fig. 2d) in IG-CA2 subtypes. However, one subtype “IG-CA2 Xpr1” (subtype 1 in Fig. 2c, 152 cells) was located in the Anterior Cingulate Area (ACA, 72 cells) and Lateral Septal Complex (LSX, 71 cells), which are anatomically distinct from the hippocampus (Extended Data Fig. 1). In-situ hybridization (ISH) data from the Allen Brain Atlas (ABA) of *Ntf3* (Fig. 2d, S4c) indicates that those cells potentially come from the IG region^32^ (Fig. 2e), which is a thin layer of gray matter lying dorsal to the corpus callosum at the base of the anterior half of the cingulate cortex, included in the ACA and LSX dissection regions (Fig. 2c, 3D model). With the power of single-cell epigenomic profiling, we are able to classify cells from this region without additional dissection. Moreover the similarity of their DNA methylomes indicates the possible functional and/or developmental relationship between the CA2 and IG excitatory neurons. Finally, the hypo-mCG regions surrounding the marker genes like *Ntf3* (Extended Data Fig. 4b) identify candidate cell-type-specific enhancers which may further our understanding of how specific gene expression programs are regulated in these structures.

**Figure 2.**
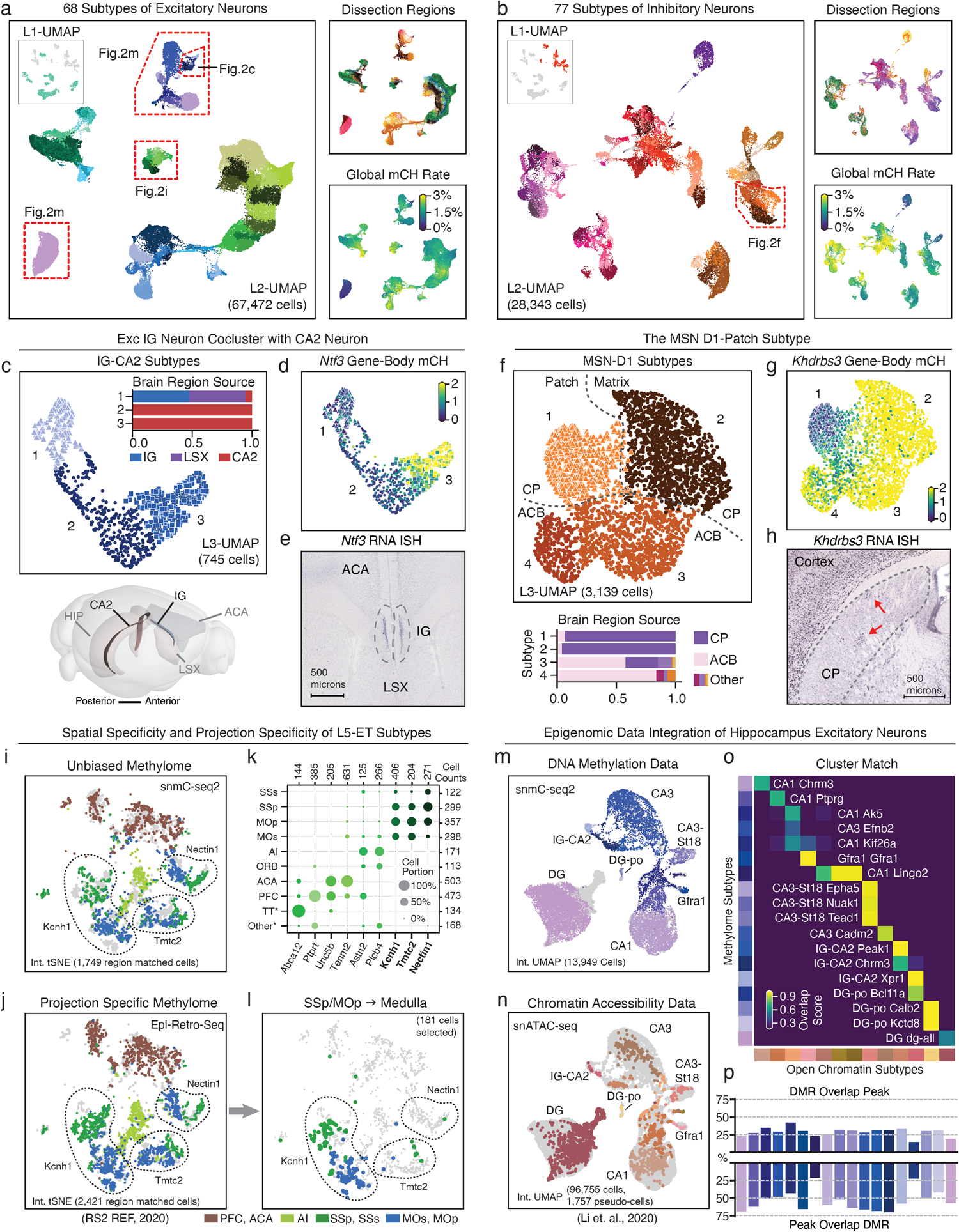
Cellular and spatial epigenomic diversity of neurons. **a, b**, Level 2 UMAP of excitatory neurons (**a**) and inhibitory neurons (**b**), colored by subtypes, dissection regions, and global mCH rate. **c**, Level 3 UMAP of IG-CA2 neurons colored by subtypes. Barplot showing sub-region composition of the subtypes: 1) “Xpr1”, 2)”Chrm3”, and 3) “Peak1”. The 3D model below illustrates these regions’ spatial relationships. **d, e**, mCH rate and an ISH experiment from the Allen brain atlas (ABA) of the *Ntf3* gene. **f**, Level 3 UMAP of MSN-D1 neurons colored by subtypes. Barplot showing sub-region composition of the subtypes: 1)”Khdrbs3”, 2) “Hrh1” 3) “Plxnc1” and 4) “Ntn1”. **g, h**, mCH rate (**g**) and an ISH experiment from ABA (**h**) of the *Khdrbs3* gene. **i, j**, Integration t-SNE on L5-ET cells profiled by snmC-seq2 (**i**) and Epi-Retro-Seq (**j**), colored by matched dissection regions in both studies. The positions of three SSp/MOp enriched subtypes are circled and labeled by their marker gene, see Extended Data Fig. 4i for the full subtype labels. **k**, L5-ET subtype spatial composition, each column sum to 100%. **l**, Medulla projecting neurons from SSp/MOp dissections profiled by Epi-Retro-Seq. **m, n**, Integration UMAP of the hippocampus excitatory neurons profiled by snmC-seq2 (**m**) and snATAC-seq (**n**, showing pseudo-cells). **o**, Overlap score matrix matching the open chromatin subtypes to the methylome subtypes. **p**, Overlap of subtype matched DMR and ATAC peaks (see Methods).

#### Methylation signatures of striatal medium spiny neurons located in patch compartments

A major GABAergic inhibitory cell type in the striatum, the *Drd1+* medium spiny neuron (MSN-D1) from the caudoputamen (CP, dorsal) and nucleus accumbens (ACB, ventral), is further separated into four subtypes (Fig. 2f, S4d). Two subtypes “MSN-D1 Plxnc1” (subtype 3 in Fig.2f) and “MSN-D1 Ntn1” (subtype 4 in Fig.2f), mainly (79%) from the ACB, are further separated by location along the anterior-posterior axis (Extended Data Fig. 4e, based on dissection), indicating spatial diversity may exist in addition to the canonical dorsal-ventral gradient of the striatum^33,34^. “MSN-D1 Khdrbs3” and “MSN-D1 Hrh1” subtypes are mostly (94%) from CP dissections, one of them marked by gene *Khdrbs3* and its potential regulatory elements (Fig. 2g, h, Extended Data Fig. 4f) corresponds to the neurochemically defined patch^35^ compartments in CP. Here the methylome profiling data provides evidence of previously unseen spatial epigenetic diversity in the striatum D1 neurons, which is also observed in the other major type MSN-D2 (*Drd2*^*+*^) of the striatum (Extended Data Fig. 4g, e, i).

#### Spatial and projection specificity of extra-telencephalic (ET) neuron subtypes

We found that cortical excitatory subtypes have distinct spatial distributions. For example, subtypes of L5-ET consist of cells from different cortical regions (Fig. 2i, k). This is further confirmed by integrating our snmC-seq2 data with data from neurons with defined projections from a parallel study using Epi-Retro-Seq (Companion Manuscript # 10)^36^ (Fig. 2j). Epi-Retro-Seq uses retrograde viral labeling to select neurons projecting to specific target brain regions followed by methylome analysis of their epigenetic subtypes. To further infer the projection target of L5-ET cells profiled in our study, we performed co-clustering on the integrated datasets and calculated the overlap between unbiased (snmC-seq2) and targeted (Epi-Retro-Seq) profiling experiments (Extended Data Fig. 4m). Interestingly, some subtypes identified from the same cortical area show different projection specificity. For example, in the L5-ET neurons, SSp and MOp neurons were mainly enriched in three subtypes marked by *Kcnh1, Tmtc2*, and *Nectin1*, respectively. However, medulla projecting neurons in the MOp and SSp only integrate with the subtype marked with *Kcnh1* (Fig. 2l), suggesting that the subtypes identified in unbiased methylome profiling have distinct projection specificity.

#### Consensus epigenomic profiles of brain cell subtypes

Integrating single-cell datasets collected using different molecular profiling modalities (e.g. snRNA-seq, snmC-seq2, snATAC-seq) can help to establish a consensus cell type atlas^23,37^. Using a mutual nearest neighbor based approach^38,39^ (see Methods), we integrated the methylome data with the chromatin accessibility data profiled using snATAC-seq on the same brain samples from a parallel study (Li et al., Companion Manuscript # 11). As visualized by the ensemble UMAP, after integration the two modalities validated each other at the level of subtype cluster assignment (Fig. 2m, n, Extended Data Fig. 5a). We then performed co-clustering of the integrated data and calculated Overlap Scores (OS) between the original methylation subtypes (m-types) and the chromatin accessibility subtypes (a-types) (Fig. 2o, Extended Data Fig. 5b). The OS describes the level of co-clustering of two groups of cells, where a higher value means the two groups are more similarly distributed across co-clusters (see Methods and Supplementary Table 7). The high overlap of CG-DMRs and open chromatin peaks in the hippocampus also confirmed the correct match of cell-type identities (Fig. 2p).

### Regulatory hierarchy of neuronal subtypes

Having developed a consensus map of cell types based on their DNA methylomes, we further explored the gene regulatory relationship between neuronal subtypes. We constructed two phylogeny trees for 68 excitatory types (Extended Data Fig. 6a) and 77 inhibitory types (Extended Data Fig. 6b) respectively, based on the gene body mCH level of 2,503 differentially methylated genes (CH-DMGs) between every pair of clusters. The taxonomy tree structures represent the similarities of these discrete subtypes, and may reflect the developmental history of neuronal type specification^22,25^.

A unique advantage of single-neuron methylome profiling is that it captures the information of both cell-type-specific gene expression and predicted regulatory events. Specifically, gene body mCH is predictive of gene expression in neurons^2,7,8^, while CG-DMRs indicate cell-type specific regulatory elements, and TFs whose motifs enriched in these CG-DMRs predict the crucial regulators of the cell type^6-8^. We used both CH-DMGs and CG-DMRs to further annotate the tree and explore the features specifying cell subtypes (Exc: Fig. 3a-c, Inh: Extended Data Fig. 6c, d).

**Figure 3.**
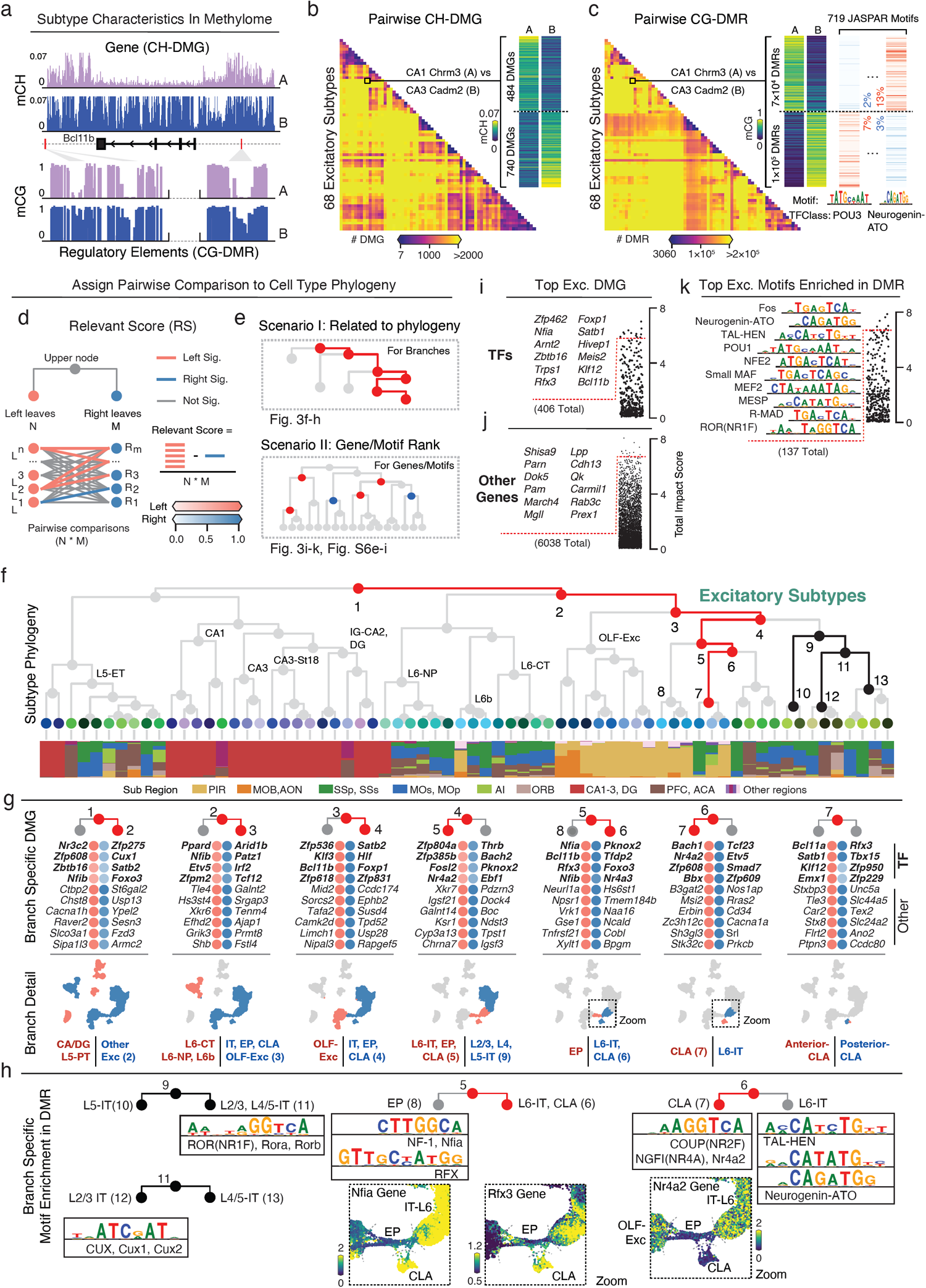
Relating genes and regulatory elements to cell subtype phylogeny. **a**, Genome browser schematic of the two characteristics contained in the methylome profiles. Genes used here are *Bcl11b*, with mCH and mCG rates from “CA1 Chrm3” (A) and “CA3 Cadm2” (B). **b, c**, Pairwise CH-DMG (**b**) or CG-DMR (**c**) counts heatmap between 68 excitatory subtypes. The detailed example from the same cluster pair as in (**a**). In each DMR set, we further scan the occurrence of 719 JASPAR motifs, two differentially enriched motifs from POU3 and Neurogenin-ATO family are shown for the example pair in (**c**). **d-e**, Schematic of impact score calculation (**d**) and two scenarios of discussing impact scores (**e**). **f**, Excitatory subtype phylogeny tree. Leaf nodes are colored by subtypes, and the barplot shows subtype composition. The numbered nodes correspond to the panels below. **g**, Top impact scores of ranked genes for the left and right branches of node 1-7. Top four TF genes are in bold, followed by other protein-coding genes. The scatter plot below shows the L2-UMAP of excitatory cells (from Fig. 2a) colored by cells involved in each branch. **h**, Branch specific TF motif families. The zoomed UMAP plots show particular TF genes in those families, whose differential mCH rate concordant with their motif enrichment. **i-k**, Total impact of TFs (**i**), TF motifs (**k**), and other protein-coding genes (**j**). Top ranked items are listed on the left.

To better understand which genes contribute to cell type specifications in the branches of the tree, we calculated a branch-specific “methylation impact score” for each gene that summarises all of the pairwise comparisons related to that branch (Fig. 3d; see Methods). The impact score ranges from 0 to 1, with a higher score predicting stronger functional relevance to the branch. With impact scores > 0.3, a total of 6,038 unique genes were assigned to branches within the excitatory phylogeny (5,975 in inhibitory), including 406 TF genes (412 in inhibitory).

Key TFs determine the neuronal cell-type identity by targeting downstream genes through the binding to regulatory DNA elements during brain development^40^. Timed expression of these TFs and their co-regulators throughout the neuron’s lifespan is critical to maintain cell-type identity^40-42^. To better understand what regulatory elements and TFs contribute to cell-type specification in the tree, we annotated branches of the tree with TF motif usage. We first identified 3.9 million DMRs across all the cell types, with an average size of 624 ± 176 bp (mean ± SD), covering 50.0% of the genome CpG sites spanning 1,240 Mb in total (see Methods). We next scanned 719 known TF binding motifs from the JASPAR database^43^ across these DMRs and performed motif enrichment analyses using all pairwise DMRs identified between neuronal subtypes (Exc: Fig. 3c, Inh: Extended Data Fig. 6d). Similar to the analysis of DMGs, the pairwise enriched motifs were assigned to branches on the subtype phylogeny using their impact scores.

This analysis revealed a possible regulatory role for several TFs assigned to the upper nodes of the phylogeny. For example, motifs from the ROR(NR1F) family were assigned to the branch that separates superficial layer IT neurons from deeper layer IT neurons (Fig. 3h, node 9), whereas motifs from the CUX family were assigned to the L2/3-IT branch separating from L4/5-IT neurons (Fig. 3h, node 11). Both families contain known members, for example *Cux1, Cux2*^*44*^, *Rorb*^*45*^ that show laminar expression in the corresponding layers and regulate cortical layer differentiation during development.

After impact score assignment, each branch of this phylogeny was associated with multiple TF genes and motifs, which potentially function in combination to shape cell-type identities^46^ (Fig. 3g, h). As an example, we focused on two brain structures of interests: the claustrum (CLA) and the Endopiriform Nucleus (EP)^47,48^. Single excitatory neurons from these two structures were included in the dissection of several cortical and piriform cortex regions (Supplementary Table 6). At the major-cell-type level, two distinct clusters are marked by *Npsr1* (EP) or *B3gat2* (CLA), and the known EP/CLA marker TF *Nr4a2*^*47,49*^ were observed to be significantly hypomethylated in both clusters compared to the others. Accordingly, the NR4A2 motif is also associated with a branch that splits CLA neurons from L6-IT neurons (Fig. 3g, h, node 6). On another branch separating EP from CLA and L6-IT neurons, several TFs including NF-1 family gene *Nfia* and *Nfib*, and RFX family gene *Rfx3*, together with corresponding motifs (Fig. 3g, 3h, node 5) rank near the top. Our findings suggest that these TFs may function together with *Nr4a2*, potentially separating EP neurons from CLA and L6-IT neurons.

Beyond identifying specific cell subtype characteristics, we hypothesized that ranking of genes or motifs by methylation variation may provide a route toward understanding their relative importance in cell type diversification and/or function. Thus, for each gene or motif, we calculated a total impact (TI) score to summarize their variation among the subtypes, using all the branch specific impact scores (see Methods). Genes or motifs with higher TI score impact more branches of the phylogeny, thus are predicted to have higher importance in distinguishing multiple subtypes. Intriguingly, by comparing the TI scores of genes and motifs calculated from the inhibitory and excitatory phylogenies, we found more TF genes and motifs having large TI scores in both cell classes than specific to either one (Extended Data Fig. 6i). For instance, *Bcl11b* distinguishes “OLF-Exc” and Isocortex IT neurons in the excitatory lineage, and distinguishes “CGE-Lamp5” and “CGE-Vip” in the inhibitory lineage. Similarly, *Satb1* separates L4-IT from L2/3-IT, and MGE from CGE in excitatory and inhibitory cells, respectively. These findings indicate broad repurposing of TFs for cell type specification among distinct developmental lineages.

In contrast, we also find TF genes and motifs only having large TI scores in one cell class (Extended Data Fig. 6i). For example, *Tshz1* gene body mCH shows a striking difference of diversity between excitatory and inhibitory cells (Extended Data Fig. 6h), suggesting that it may function in shaping inhibitory subtypes, but not excitatory subtypes. Similarly, bHLH DNA binding motifs show much higher TI scores for excitatory subtypes compared with inhibitory (Extended Data Fig. 6i). While genes in this TF family such as *Neurod1/2* have long been known to participate in excitatory neuron development, they have not been reported to regulate GABAergic neuron differentiation^50^.

### Defining regulatory interaction between enhancers and genes at the cell type level

To systematically identify enhancer-like regions in specific cell types, we predicted enhancer-DMRs (eDMR) by integrating matched DNA methylome and chromatin accessibility profiles of 161 subtypes using the REPTILE algorithm^51^ (Fig. 4a). We identified 1,612,198 eDMR (34% of total DMRs), 73% of which overlapped with separately identified snATAC peaks (Fig. 4b). Fetal-enhancer DMRs (feDMR, eDMRs identified across developing time points) of forebrain bulk tissues^6^ show high (88%) overlap with eDMRs. Surprisingly, the eDMRs also cover 74% of the feDMRs from other fetal tissues^6^, indicating extensive reuse of enhancer-like regulatory elements across mammalian tissue types (Fig. 4b).

**Figure 4.**
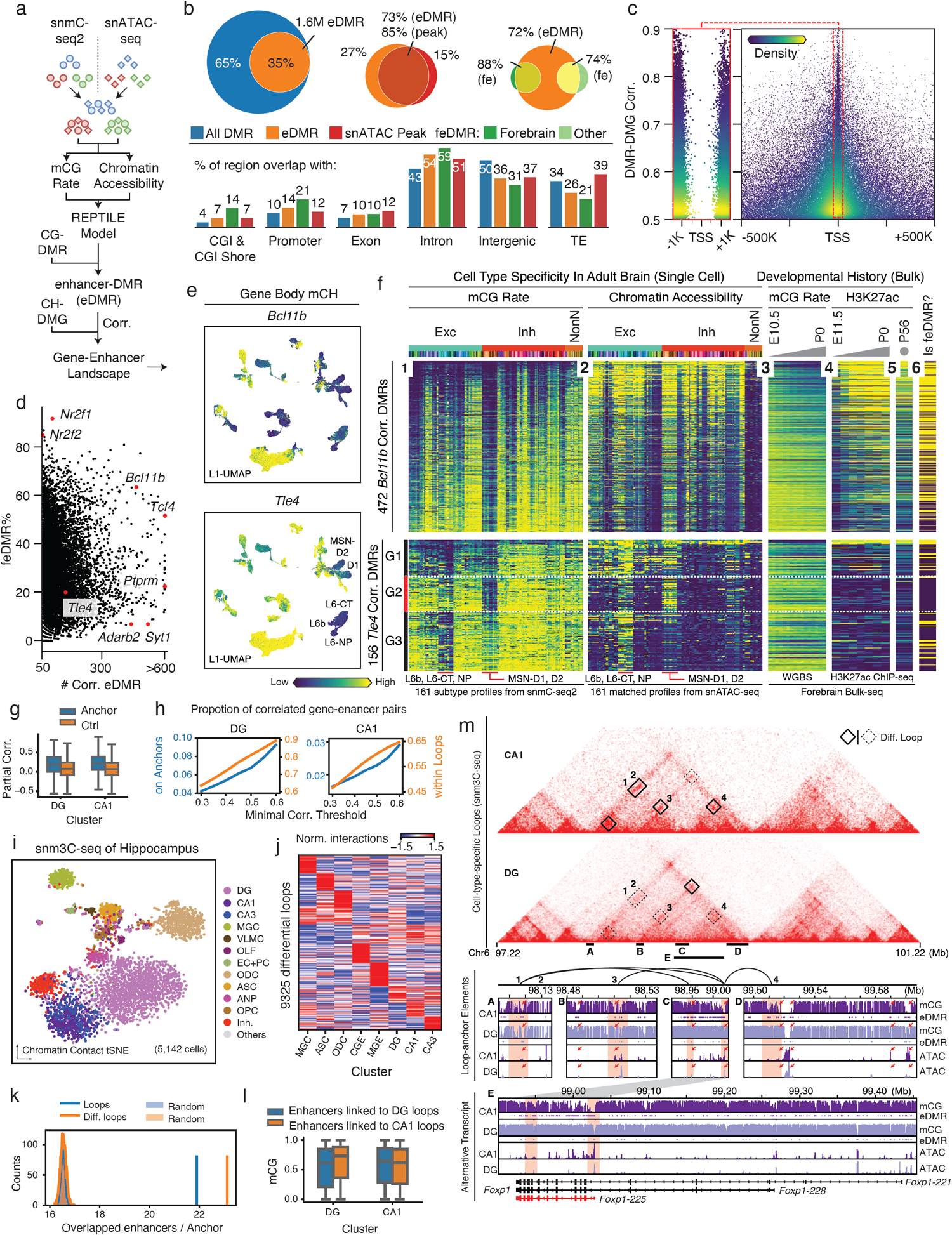
Gene-enhancer landscapes in neuronal subtypes. **a**, Schematic of enhancer calling using matched DNA methylome and chromatin accessibility subtype profiles. The REPTILE algorithm was deployed followed by building the gene-enhancer landscape through their methylation correlation. **b**, Overlap of regulatory elements identified in this study, and other epigenomic studies (snATAC-seq peaks from Li. et al, Companion Manuscript 11; fetal enhancer DMRs^6^). **c**, DMR-DMG correlation (y-axis) and the distance between DMR center and gene TSS (x-axis), each point is a DMR-DMG pair colored by kernel density. **d**, Percentage of positively correlated eDMR that overlap with forebrain feDMR (from He et al.^6^). Each point is a gene, while x-axis is the number of positively correlated eDMRs to each gene. **e**, The gene body mCH rate of *Bcl11b* (top) and *Tle4* (bottom) gene. **f**, The predicted enhancer landscape of *Bcl11b* (top) and *Tle4* (bottom). Each row is a correlated eDMR to the gene, columns from left to right are: (1) mCG rate and (2) ATAC FPKM in 161 subtypes; (3) bulk developing forebrain tissue mCG rate^6^ and (4) H3K27ac FPKM^68^; (5) adult frontal cortex H3K27ac^2^; and (6) is a feDMR or not^6^. **g**, Partial correlation between mCG of enhancers and mCH of genes on separated loop anchors of DG (left) and CA1 (right) compared to random anchors with comparable distance (n=4,171, 4,036, 4,326, 5,133). **h**, Proportion of loop supported enhancer-gene pairs among those linked by correlation analyses surpassing different correlation thresholds. **i**, t-SNE of sn-m3C-seq cells (n=5,142) colored by clusters. Cluster labels were predicted using the model trained with snmC-Seq data, and some major cell-types were merged. **j**, Interaction level of 9,325 differential loops in eight clusters at 25kb resolution. Values shown are z-score normalized across rows and columns. **k**, Number of enhancers per loop anchor (blue) or per differential loop anchor (orange) compared to randomly selected 25kb regions across the genome. The permutation was repeated for 2,000 times. **l**, mCG of enhancers linking to DG specific loops (blue, n=13,854) and CA1 specific loops (orange, n=14,373) in DG (left) or CA1 (right). **m**, Comprehensive epigenomic signatures surrounding *Foxp1*. The triangle heatmap shows CA1 and DG chromatin contacts with differential loops labeled with black boxes; the next genome browser section shows detail mCG and ATAC profiles near four CA1-specific loops’ anchors, with red rectangles indicate loop anchor and red arrows indicate notable regulatory elements; genome browser image depicts the mCG and ATAC profiles at the *Foxp1* gene. The elements of boxplots in (**g**) and (**l**) are defined as: center line, median; box limits, first and third quartiles; whiskers, 1.5x interquartile range.

Next, we examined the relationship between the cell type signature genes and their potential regulatory elements. After regressing out the global methylation level, we calculated the partial correlation between all DMG-DMR pairs within 1 Mb distance, using methylation levels across 145 neuronal subtypes (see Methods). Non-neuronal subtypes were not included in this analysis due to the large difference in the global methylation level compared to neurons. We identified a total of 1,038,853 (64%) eDMR that correlated with at least one gene (correlation > 0.3 with empirical *P* value < 1e-4, two-sided test, Extended Data Fig. 7a). Notably, for those strongly positive-correlated DMR-DMG pairs (correlation > 0.5), the DMRs are largely (63%) within 100kb to the TSSs of the corresponding genes, but depleted from ± 1kb (Fig. 4c, Extended Data Fig. 7b), whereas for the negatively correlated DMR-DMG pairs, only 11% of DMRs are found within 100kb of the TSS (Extended Data Fig. 7c).

eDMR-DMG correlation analysis predicts regulatory interactions between enhancers and genes at subtype resolution. Using the gene-enhancer interactions predicted by this correlation analysis, we assigned eDMRs to their target genes, and calculated the percentage of overlap with feDMR. We found that these percentages vary dramatically among genes (Fig. 4d). For example, in *Bcl11b*, an early developmental TF gene^52^, 63% of the positively correlated eDMRs overlap with forebrain feDMRs (Fig. 4f, first row). However, for *Tle4*, a gene known to be functional in L6-CT^53^ and MSN-D1/D2 neurons, the percentage of eDMR that overlapped with feDMRs was only 20% (Fig. 4d). Interestingly, when further classifying *Tle4* correlated eDMR into three subgroups using K-means clustering (k=3, see Methods), we identified subgroups with different cell-type-specificity (Fig. 4f, second row. Extended Data Fig. 7d). One group (G2) of elements that displayed little diversity during development in bulk data showed highly specific mCG and open chromatin signals in MSN-D1/D2 neurons, while another group (G3) is specific to L6-CT neurons. These two groups of DMRs suggest possible alternative regulatory elements usage to regulate the same gene in different cell-types although further experiments are required to validate this hypothesis.

Together, these analyses indicate that previously defined developmental regulatory elements from bulk tissues are limited in terms of the cell type resolution. For those broadly effective regulatory elements (e.g., feDMRs correlated with *Bcl11b*) that also have been identified in bulk tissues, single-cell profiling allows us to carefully chart their cell type specificity into subtype level. In addition, we identified many novel regulatory elements (e.g., eDMRs correlated with *Tle4* in MSN-D1/D2) that show more restricted specificity, providing abundant candidates for further pursuing enhancer-driven AAVs^54-56^ that target fine cell types.

### Single-nucleus multi-omic profiling of chromatin contact and DNA methylation validates gene-enhancer interactions

Distal enhancers typically regulate gene expression by physical interaction with promoters^57^. Therefore, to examine whether our correlation-based predictions of enhancer-gene associations are supported by physical chromatin contacts, we generated single-nucleus methylation and chromosome conformation capture sequencing (sn-m3C-seq)^18^ data for 5,142 single nuclei from the DG and CA (152k contacts per cell on average). We assigned each of these cells to one of the 161 subtypes based on their methylome, using a supervised model trained with >100k snmC-Seq2 cells. By merging the contact matrices of all single cells belonging to the same methylation-defined subtype, we generated contact maps for eight major cell types that include more than 100 cells. In total, 19,151 chromosome loops were identified in at least one of the cell-types at 25kb resolution (range from 1,173 to 12,614).

Since sn-m3C-seq data identified structural interactions between genes and regulatory elements, we next asked whether these interactions were consistent with the predictions from eDMR-DMG correlation analysis. Using DG and CA1 as examples (the neuronal types with highest coverage), a notably higher correlation was observed between enhancers and genes at loop anchors compared to random enhancer-gene pairs after controlling for genomic distance (Fig. 4g; *P* value=5.9e-74 for DG and 3.0e-158 for CA1, two-sided Wilcoxon rank-sum tests). Reciprocally, the enhancer-gene pairs showing the stronger correlation of methylation were more likely to be found linked by chromosome loops or within the same looping region (Fig. 4h, Extended Data Fig. 7f). We also compared the concordance of methylation patterns between genes and enhancers linked by different interaction-based methods, and found the pairs linked by loop anchors or closest genes had the highest correlation of methylation (Extended Data Fig. 7g). Together, these analyses validate the physical proximity of co-associated, hypomethylated enhancer-gene pairs predicted by our correlation based method in specific cell types.

In addition, we observed significant cell type specificity of 3D genome structures. The major cell types could be distinguished on UMAP embedding based on chromosome interaction at 1 Mb resolution (Fig. 4i), indicating the dynamic nature of genome architecture across cell types. Among the 19,151 chromosome loops at 25 kb resolution, 48.7% showed significantly different contact frequency between cell types (Fig. 4j). eDMRs were highly enriched at these differential loop anchors (Fig. 4k; p<0.005, permutation test). mCG levels at distal cis-elements are typically anti-correlated with enhancer activity^6^. Thus, we hypothesized that enhancers at differential loop anchors might also be hypomethylated in the corresponding cell-type. Indeed, using the loops identified in DG granule cells and CA1 as an example, we observed that in DG, enhancers at the anchors of DG specific loops were hypomethylated compared to enhancers at the anchors of CA1 specific loops (*P* value=2.9e-103, two-sided Wilcoxon rank-sum test); the opposite scenario was found in CA1 (Fig. 4l; *P* value=3.5e-5, two-sided Wilcoxon rank-sum test).

Many differential loops were observed near the marker genes of the corresponding cell type. For example, *Foxp1*, a highly expressed TF in CA1 but not DG^58^, has chromosome loops surrounding its gene body in CA1 but not DG (Fig. 4m). eDMRs and open chromatin were observed at these loop anchors (Fig. 4m). Intriguingly, three loops in CA1 anchored at the TSS of the same transcript of *Foxp1* (Fig. 4m, box 2-4). Stronger demethylation and chromatin accessibility were also observed at the same transcript compared to the others (Fig. 4m, view E). These epigenetic patterns might suggest a specific transcript of *Foxp1* (Foxp1-225) is selectively activated in CA1. In contrast, *Lrrtm4*, a DG marker gene encoding a presynaptic protein that mediates excitatory synapse development in granule cells^59^, shows extensive looping to distal elements in DG but not CA1 (Extended Data Fig. 7h). Notably, 34 genes showed alternative loop usage, 20 of which expressed in both DG and CA1 (CPM > 1). *Grm7* is an example where its TSS interacts with an upstream enhancer in DG but with gene-body enhancers in CA1 (Extended Data Fig. 7i).

### Spatial gradients in intra-telencephalic (IT) excitatory neurons

IT neurons from different cortical areas are typically classified into major types corresponding to their laminar locations: L2/3, L4, L5, and L6 (Fig. 5a). In agreement with the correlation between transcription^22^ and DNA methylation, we found that IT neurons show hypomethylation of the marker genes defined in Tasic et al.^22^ in the corresponding layers (Extended Data Fig. 8c). For example, the *Rorb*^+^ L4 cells show unique gene body hypomethylation in both somatomotor (MO) and somatosensory areas (SS) (Extended Data Fig. 8c). These findings suggest that although the MO lacks a cytoarchitectural visible L4^60^, there is a population of IT neurons that are epigenetically similar to L4 neurons in SS. The presence of a putative L4 neuron population in MO is also supported by transcriptomics^23^ and neuronal connectivity^60^. Furthermore, unsupervised UMAP embedding (Fig. 5a) reveals a continuous gradient of IT neurons resembling the medial-lateral distribution of the cortical areas (Fig. 5b), strongly suggesting that the cortical arealization information is well preserved in the DNA methylome.

**Figure 5.**
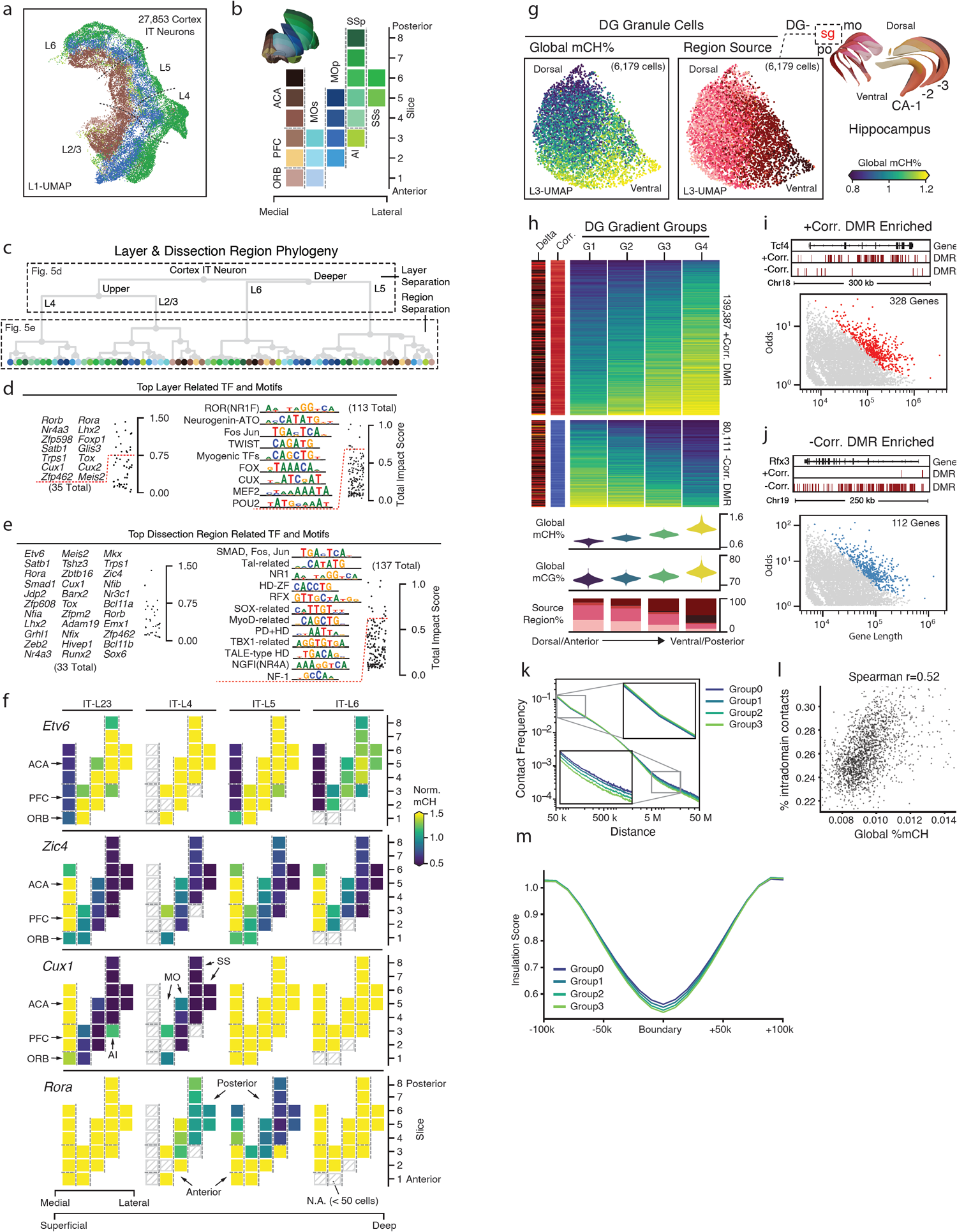
Brain-wide spatial gradients of DNA methylation. **a**, L1-UMAP for cortex IT neurons colored by dissection regions. **b**, 2-D layout of twenty-one dissection regions profiled in this study, colored by dissection regions. **c**, IT spatial group phylogeny. The top three nodes separate four layers, and downstream nodes separate dissection regions (see Extended Data Fig. 8a for details). **d, e**, The top layer (**d**) and dissection region (**e**) related TFs and JASPAR motifs ranked by total impact score (see Methods). **f**, Gene body mCH rate of TF genes in (**e**) using the same layout as (**b**). **g**, L3-UMAPs for DG granule cells colored by cell global mCH rate and dissection regions. **h**, Compound figure showing four DG gradient cell groups and the two groups of gradient DMR separated by their sign of correlation to the cell’s global mCH level. **i, j**, Gradient DMR enriched genes. Red dots indicate genes with positive (**i**) or negative (**j**) DMR enriched in their gene body. DMRs of *Tcf4* and *Rfx3* gene. Genome browser view of representative positive and negative correlated DMR enriched genes respectively. **k**, Interaction frequency decays with increasing genome distances in different groups. **l**, Correlation between global mCH and proportion of intra-domain contacts across 1,904 DG cells. **m**, Insulation scores of 9,160 domain boundaries and flanking 100kb regions.

To systematically explore the spatial gradient of DNA methylation patterns, we remerge the cells into spatial groups based on their cortical layer and anatomical regions. A phylogenetic tree was generated using the mCH level of highly variable 100kb chromosome bins of these spatial groups’ centroids (see Methods). The phylogeny split the cells into four laminar layer groups, followed by cortical area separation within each layer (Fig. 5c). This provides a clear structure for calculating the methylation total impact score of layer-related or region-related DMGs and TF binding motifs separately (Fig. 5d, see Methods). In brief, the top layer-related TFs included most of the known laminar marker genes together with their DNA binding motifs (Fig. 5d), while some of them show regional specific methylation differences between layers. For example, *Cux1*, a top-ranked TF marker gene for L2/3 and L4 neurons, is hypomethylated in MO and SS, but is hypermethylated in L2/3 of other regions we sampled, in agreement with patterns from in-situ hybridization^61^. *Cux2*, a gene from the same TF family, does not show the same regional specificity (Extended Data Fig. 8c). We also identified many additional TFs having cortical region specificity (Fig. 5e, f). For example, *Etv6* is only hypomethylated in medial dissection regions, while *Zic4* is hypermethylated in those regions across layers. In contrast, *Rora* shows anterior-posterior methylation gradient only in the L4 and L5 cells. Together, these observed methylome spatial gradients demonstrated the value of our dataset for further exploring the cortical arealization with cell-type resolution.

### Spatial gradients in dentate gyrus granule cells

Another striking spatial gradient was observed in granule cells from DG. UMAP embedding of granule cells displayed continuous global mCH and mCG level gradients that correlated with their dorsal-ventral location. Granule cells from the ventral DG show higher global CG and CH methylation compared to cells from the dorsal DG (Fig. 5g). To investigate the biological significance of these gradients, we identified 219,498 gradient CG-DMRs by grouping DG cells according to their global mCH level. Among them, 139,387 DMRs are positively correlated with global mCH, and 80,111 are negatively correlated DMRs (Fig. 5h). Positively or negatively correlated DMRs showed significant enrichment in two different sets of genes (245 pos. corr.; 183 neg. corr.) which include TF genes such as *Tcf4* and *Rfx3* compared to the genome background (Fig. 5i-j, see Methods). Based on Gene Ontology analysis, the negatively correlated DMRs are enriched in genes related to synaptic functions (Extended Data Fig. 8d), while the positively correlated DMRs are enriched in developmental related genes (Extended Data Fig. 8e). Together, these results indicate a continuous methylation gradient exists in the dorsal-ventral axis of the DG granule layers, supporting an intra-cell-type molecular shift that parallels the known functional and anatomical differences in dorsal-ventral DG^62,63^ that are possibly controlled via gradient DNA methylation^64^.

Next, we investigated whether the global dynamic of methylation is correlated with changes in 3D genome architecture. By plotting the interaction strength against the genomic distance between the anchors (decay curve), a higher proportion of short-range contacts and smaller proportion of long-range contacts were observed in the groups with higher global mCH (Fig. 5k). This might indicate a more compact nuclear structure in these groups. The genome is organized into specific 3D features with different levels of resolution, including compartments, domains, and loops. We tested the differences of these features across DG cells grouped based on their global mCH levels. Although compartment strengths were not correlated with these methylation changes (Extended Data Fig. 8f), the number of intra-domain contacts was positively correlated with global mCH across single cells (Fig. 5l), which is concordant with the patterns observed in the contact frequency decay curves. Interestingly, after normalizing the effect of decay, we found the insulation scores at domain boundaries were significantly lower in the groups with high global mCH levels (Fig. 5m; all p<1e-10, two-sided Wilcoxon signed-rank test), which suggests that the domain structures might be more condensed over flanking regions in these cell groups.

### Deep neural network learning of cell identity and spatial location

To further test the extent that spatial and cell-type information is encoded in a single neuron’s DNA methylome, we built a multi-task deep Artificial Neural Network (ANN) using cell-level methylome profiles in this study (Fig. 6a). Specifically, mCH rates of non-overlapping 100kb bins from 94,383 neurons were used to train and test the network with five-fold cross-validation. The ANN was able to predict subtype identity and spatial location simultaneously for each testing cell with 95% and 89% accuracies, respectively (Fig. 6b-d). Importantly, the location prediction accuracy using DNA methylation is significantly higher than only using the spatial distribution information of each subtype (overall accuracy increased by 38%, Extended Data Fig. 9b), suggesting that spatial diversity is well-preserved in the neuronal DNA methylome. We did, however, notice errors in location prediction in a few cell types, most notably in MGE and CGE derived inhibitory neurons (Fig. 6e, Extended Data Fig. 9b). This finding is consistent with previous transcriptome based studies^22^, suggesting these inhibitory neurons do not display strong cortical region specificity.

**Figure 6.**
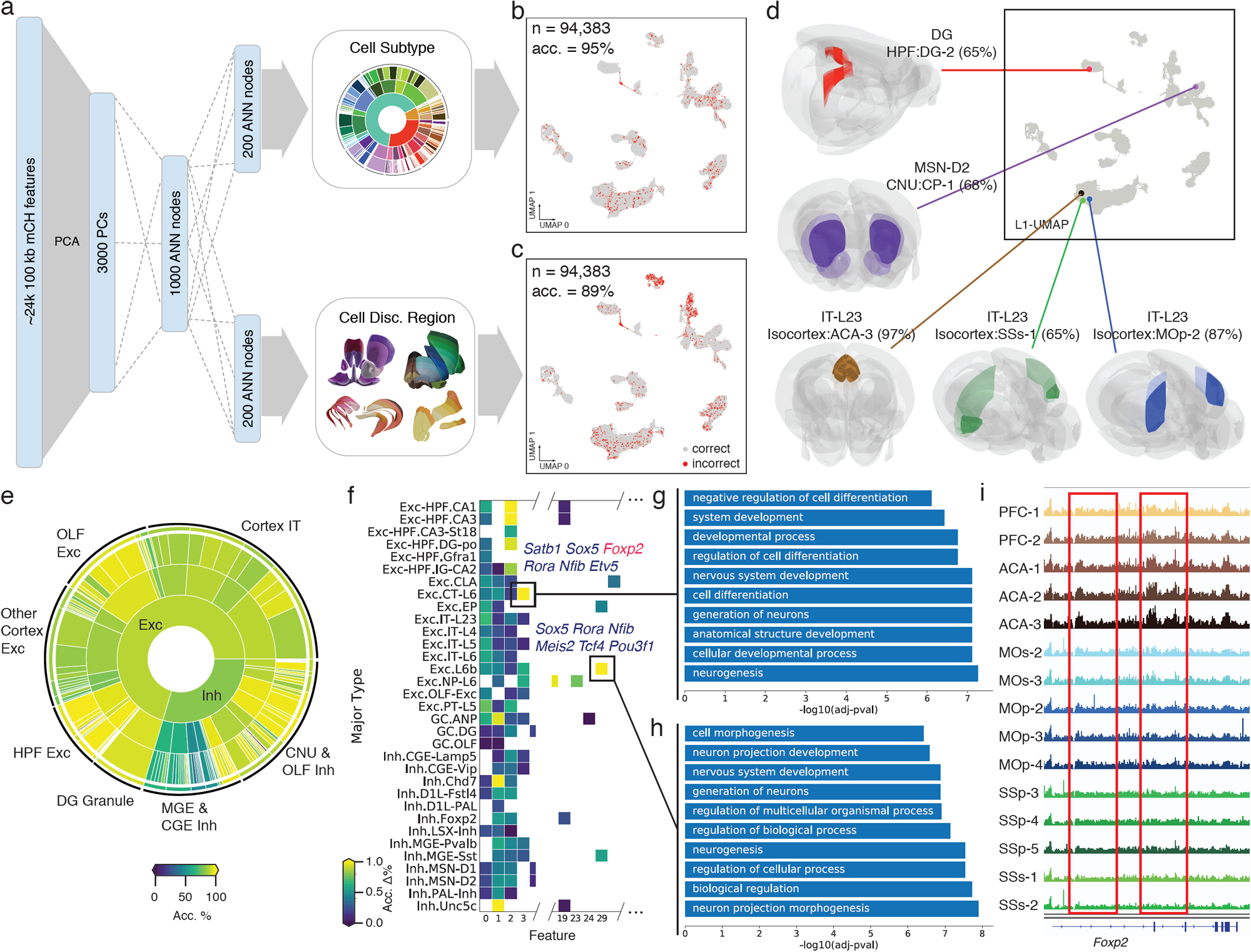
A methylome-based predictive model captures both cellular and spatial characteristics of neurons. **a**, The architecture of the artificial neural network for predicting both cell-type identity and spatial origin. **b, c**, Performance of subtype (**b**) and dissection region (**c**) predictions. incorrect predictions are colored in red on L1 UMAP. **d**, Examples of using the model to predict cell type identity and spatial location simultaneously. The color intensity denotes the probability that the cell came from the discretion region. The highest probabilities are shown in brackets. **e**, The overall predictability of spatial location for each level of neuronal types (see Extended Data Fig. 9b for details). **f**, Feature importance evaluation for spatial origin prediction. **g, h**, GO term enrichment of top-loading genes of features that are important for predicting the spatial location of L6-CT (**g**) and L6b (**h**). **i**, Genome browser view of the mCH rate of the *Foxp2* gene in each cortical dissection region.

Next, we selected the features (100kb bins) that capture most spatial information (Fig. 6f, see Methods), and found that genes related to neuronal system development are highly enriched in these bins (Fig. 6g, h). Many cell type marker genes are also located in these bins, such as *Foxp2* (Fig. 6f). In addition to distinguishing L6-CT neurons from other cell types, *Foxp2* shows notably mCH difference among different dissected regions in L6-CT (Fig. 6I), indicating the extensive impact of neuron spatial origin on their methylomes.

## Discussion

In this study, we describe the generation and analysis of a comprehensive single-cell DNA methylation atlas for the mouse brain, encompassing over 110,000 methylomes; the largest methylation dataset for any organism to date. By profiling of 45 brain regions, a total of 161 subtypes: 68 excitatory, 77 inhibitory, and 16 non-neuronal subtypes were classified using the three-levels of iterative clustering. These subtypes are associated with a total of 24,208 CH-DMGs and 3.9M CG-DMRs, covering 50% (1,240 Mb) of the mouse genome. Through integration with snATAC-seq (Li et al. Companion Manuscript # 11), we matched the chromatin accessibility clusters to each of the methylome subtypes and used the combined epigenomic information to predict 1.5M active-enhancer-like eDMRs, including 72% cell-type-specific elements that were missed from previous tissue level bulk studies^6,65^.

To describe cell-type specificity of genes and TF binding motifs in the context of cell type phylogeny, we defined a metric called the methylation impact score. This metric aggregates pairwise DMGs and DMRs between subtypes, and assigns them to branches of a cell type phylogeny. These assignments allow us to describe cell-type specificity at different levels of the phylogeny, potentially relating to different stages of their neuronal developmental lineages. We found that many known TF genes and their corresponding DNA-binding motifs were co-associated with the same branch in the phylogeny, supporting the biological significance of this approach. An important outcome of this analysis is the discovery of many novel TFs with high impact scores, providing a rich source of candidate TFs for future study.

To study the associations of eDMRs and their targeting genes, we first established the eDMR-gene landscape via correlation between the mCH rate of genes and mCG rate of DMRs. Furthermore, in the hippocampus, we also generated high-resolution single-cell 3D chromatin conformation and DNA methylome from the same cells using sn-m3C-seq. With this multi-omic dataset, we obtained cell-type-specific loops in eight cell types. The physical loops and eDMR-gene correlation loops together linked candidate enhancers to their target genes in specific cell types.

Through the analyses of single-cell methylome from detailed spatial brain region dissections, our brain-wide epigenomic dataset reveals extraordinary spatial diversity encoded in the DNA methylomes of neurons. The iterative clustering analysis separates fine-grained spatial information, which is accurately reproduced by the ANN trained on the single-cell methylome profiles. Moreover, the ANN also reveals additional spatial specifications within most of the subtypes, indicating continuous spatial DNA methylation gradients widely exist among cell types. The spatial gradients observed for excitatory IT neurons within distinct cortical areas are of special interest. During cortex development, glutamatergic neurons are regionalized by a proto map formed from an early developmental gradient of TF expression^66,67^. Similarly we observed that many TF genes and their corresponding DNA binding motifs showed gradients of DNA methylation in adult IT neurons from distinct cortical regions. Additionally, we found intra-subtype methylation gradients in DG granule cells. These CG-DMR are enriched in essential synaptic genes, suggesting the existence of a methylation gradient in the dorsal-ventral axis in the DG granule layer^64^.

Overall, we present the first single-cell resolution DNA methylomic mouse brain atlas with detailed spatial dissection and subtype level classification. Our analysis highlights the power of this dataset for characterizing cell types with both gene activity information in the coding regions and the regulatory elements in the non-coding region. This comprehensive epigenomic dataset provides a valuable resource for answering fundamental questions about gene regulation in specifying cell type spatial diversity and provides the raw material to develop new genetic tools for targeting specific cell types and functional testing of them.

## Acknowledgments

We thank Dr. Yupeng He for the advice on the methylpy and REPTILE analysis. We thank Dr. Terrence Sejnowski for the advice on the ANN analysis. This work is supported by NIMH U19MH11483 to J.R.E. and E.M.C. The Flow Cytometry Core Facility of the Salk Institute is supported by funding from NIH-NCI CCSG: P30 014195. J.R.E is an investigator of the Howard Hughes Medical Institute.

## Author Contributions

J.R.E., H.L., B.R., M.M.B., C.L., J.R.D. conceived the study. H.L., J.Z., W.T. analyzed the snmC-seq data and drafted the manuscript. J.R.E., C.L., E.A.M., J.R.D., M.M.B. edited the manuscript. J.R.E., H.L., M.M.B., A.B., J.L., S.P. coordinated the research. M.M.B., A.B., A.A., H.L., J.L., J.R.N., A.R., J.K.O., C.O., L.B., C.F., C.L., J.R.E. generated the snmC-seq2 data. J.R.D., B.C., A.B., J.L., J.Z., A.A., J.K.O., C.L., J.R.N., C.O., L.B., C.F., R.G.C., M.M.B., J.R.E. generated the sn-m3C-seq data. S.P., M.M.B., X.H., J.L., O.B.P., Y.E.L., J.K.O., B.R. generated the snATAC-seq data. Z.Z., J.Z., E.M.C., M.M.B., J.R.E., A.B., A.A., J.R.N., C.O., L.B., C.F., R.G.C., A.R. generated the Epi-Retro-Seq data. H.L., H.C., E.A.M., M.N., C.L. contributed to data archive/infrastructure. J.R.E. supervised the study.

## Methods

### Mouse brain tissues

All experimental procedures using live animals were approved by the Salk Institute Animal Care and Use Committee under protocol number 18-00006. Adult (P56) C57BL/6J male mice were purchased from Jackson Laboratories and maintained in the Salk animal barrier facility on 12 h dark-light cycles with food ad-libitum for a maximum of 10 days. Brains were extracted and sliced coronally at 600 µm from the frontal pole across the whole brain (for a total of 18 slices) in ice-cold dissection buffer containing 2.5mM KCl, 0.5mM CaCl_2_, 7mM MgCl_2_, 1.25mM NaH_2_PO_4_, 110mM sucrose, 10mM glucose, and 25mM NaHCO_3_. The solution was kept ice-cold and bubbled with 95% O_2_ / 5% CO_2_ for at least 15 min before starting the slicing procedure. Slices were kept in 12-well plates containing ice-cold dissection buffers (for a maximum of 20 min) until dissection aided by an SZX16 Olympus microscope equipped with an SDF PLAPO 1XPF objective. Each brain region was dissected from slices along the anterior-posterior axis according to the Allen Brain reference Atlas CCFv3^69^ (see Extended Data Fig. 1 for the depiction of a posterior view of each coronal slice). Slices were kept in ice-cold dissection media during dissection and immediately frozen in dry ice for posterior pooling and nuclei production. For nuclei isolation, each dissected region was pooled from 6-30 animals, and two biological replicas were processed for each slice.

### Nuclei isolation and Fluorescence Activated Nuclei Sorting (FANS)

Nuclei were isolated as previously described^2,8^. Isolated nuclei were labeled by incubation with 1:1000 dilution of AlexaFluor488-conjugated anti-NeuN antibody (MAB377X, Millipore) and a 1:1000 dilution of Hoechst 33342 at 4°C for 1 hour with continuous shaking. Fluorescence-Activated Nuclei Sorting (FANS) of single nuclei was performed using a BD Influx sorter with an 85µm nozzle at 22.5 PSI sheath pressure. Single nuclei were sorted into each well of a 384-well plate preloaded with 2 µl of Proteinase K digestion buffer (1µl M-Digestion Buffer, 0.1µl 20 µg/µl Proteinase K and 0.9µl H2O). The alignment of the receiving 384-well plate was performed by sorting sheath flow into wells of an empty plate and making adjustments based on the liquid drop position. Single-cell (1 drop single) mode was selected to ensure the stringency of sorting. For each 384-well plate, columns 1-22 were sorted with NeuN+ (488+) gate, and column 23-24 with NeuN-(488-) gate, reaching an 11:1 ratio of NeuN+ to NeuN-nuclei.

### Library preparation and Illumina sequencing

Detailed methods for bisulfite conversion and library preparation were previously described for snmC-seq2^8,16^. The snmC-seq2 and sn-m3C-seq (see below) libraries generated from mouse brain tissues were sequenced using an Illumina Novaseq 6000 instrument with S4 flowcells and 150 bp paired-end mode.

### The sn-m3C-seq specific steps of library preparation

Single-nucleus methyl-3C sequencing (sn-m3C-seq) was performed as previously described^18^. In brief, the same batch of dissected tissue samples from the dorsal dentate gyrus (DG-1 and DG-2, see Supplementary Table 2), ventral dentate gyrus (DG-3 and DG-4), dorsal hippocampus (CA-1 and CA-2), and ventral hippocampus (CA-3 and CA-4), were frozen in liquid nitrogen. The samples were then pulverized while frozen using a mortar and pestle, and then immediately fixed with 2% formaldehyde in DPBS for 10 minutes. The samples were quenched with 0.2M Glycine and stored at −80C until ready for further processing. After isolating nuclei as previously described^18^, nuclei were digested overnight with NlaIII and ligated for 4 hours. Nuclei were then stained with Hoechst 33342 and filtered through a 0.2µM filter and sorted similarly to the snmC-seq2 samples. Libraries were generated using the snmC-seq2 method.

### Mouse brain region nomenclature

The mouse brain dissection and naming of anatomical structures in this study followed the Allen Mouse Brain Atlas^69^. Based on the hierarchical structure of the Allen CCF, we used a three-level spatial region organization to facilitate description: a. The major region, e.g., isocortex, hippocampus, b. The sub-region, e.g., MOp, SSp, within isocortex, c. The dissection region, e.g., MOp-1, MOp-2, within MOp. Supplementary Table 1 contains full names of all abbreviations used in this study.

### Analysis stages

The following method sections were divided into three stages. The first stage describes mapping and generating files in the single-cell methylation-specific data format. The second stage describes clustering, identifying differentially methylated genes (DMG), or integrating other datasets, which all happened at the single-cell level. The third stage describes the identification of putative cell-type-specific regulatory elements using cluster-merged methylomes. Other figure-specific analysis topics may combine results from more than one stage.

## STAGE I. MAPPING AND FEATURE GENERATION

### Mapping and feature count pipeline

We implemented a versatile mapping pipeline (<u>cemba-data.rtfd.io</u>) for all the single-cell methylome based technologies developed by our group^8,15,16^. The main steps of this pipeline included: 1) Demultiplexing FASTQ files into single-cell; 2) Reads level QC; 3) Mapping; 4) BAM file processing and QC; 5) final molecular profile generation. The details of the five steps for snmC-seq2 were previously described^16^. We mapped all of the reads to the mouse mm10 genome. After mapping, we calculated the methylcytosine counts and total cytosine counts for two sets of genomic regions in each cell. Non-overlapping chromosome 100kb bins of the mm10 genome (generated by “bedtools makewindows -w 100000”), were used for clustering analysis and ANN model training, and the gene body region ± 2kb defined by the mouse GENCODE vm22, were s used for cluster annotation and integration with other modalities.

### sn-m3C-seq specific steps or read mapping and chromatin contact analysis

Methylome sequencing reads were mapped following the TAURUS-MH pipeline as previously described^18^. Specifically, reads were trimmed for Illumina adaptors and then an additional 10bps was trimmed on both sides. Then R1 and R2 reads were mapped separately to the mm10 genome using bismark with bowtie. The unmapped reads were collected and split into shorter reads representing the first 40bps, the last 40bps, and the middle part of the original reads (if read length > 80bp after trimming). The split reads were mapped again using Bismark with the Bowtie. The reads with MAPQ<10 were removed. To generate the methylation data, the filtered bam files from split and unsplit R1 and R2 reads were deduplicated with picard and merged into a single bam file. Methylpy (v1.4.2)^70^ was used to generate an allc file from the bam file for every single cell. To generate the Hi-C contact map, we paired the R1 and R2 bam files where each read pair represents a potential contact. For generating contact files, read pairs where the two ends mapped within 1kbp of each other were removed.

## STAGE II. CLUSTERING RELATED

### Single-cell methylome data quality control and preprocessing

#### Cell filtering

We filtered the cells based on these main mapping metrics: 1) mCCC rate < 0.03. mCCC rate reliably estimates the upper bound of bisulfite non-conversion rate^8^, 2) overall mCG rate > 0.5, 3) overall mCH rate < 0.2, 4) total final reads > 500,000, 5) bismark mapping rate > 0.5. Other metrics such as genome coverage, PCR duplicates rate, index ratio were also generated and evaluated during filtering. However, after removing outliers with the main metrics 1-5, few additional outliers can be found.

#### Feature filtering

100kb genomic bin features were filtered by removing bins with mean total cytosine base calls < 250 or > 3000. Regions that overlap with the ENCODE blacklist^71^ were also excluded from further analysis.

#### Computation and normalization of the methylation rate

For CG and CH methylation, the computation of methylation rate from the methyl-cytosine and total cytosine matrices contains two steps: 1) prior estimation for the beta-binomial distribution and 2) posterior rate calculation and normalization per cell.

Step 1, for each cell we calculated the sample mean, *m*, and variance, *v*, of the raw mc rate *(mc / cov)* for each sequence context (CG, CH). The shape parameters (α, β) of the beta distribution were then estimated using the method of moments:

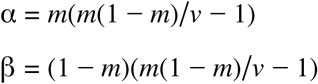

This approach used different priors for different methylation types for each cell and used weaker prior to cells with more information (higher raw variance).

Step 2, We then calculated the posterior: 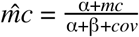., We normalized this rate by the cell’s global mean methylation, *m* = α/ (α + β). Thus, all the posterior 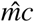 with 0 *cov* will be constant 1 after normalization. The resulting normalized *mc* rate matrix contains no NA (not available) value, and features with less *cov* tend to have a mean value close to 1.

#### Selection of highly variable features

Highly variable methylation features were selected based on a modified approach using the scanpy package *scanpy*.*pp*.*highly_variable_genes* function^72^. In brief, the *scanpy*.*pp*.*highly_variable_genes* function normalized the dispersion of a gene by scaling with the mean and standard deviation of the dispersions for genes falling into a given bin for mean expression of genes. In our modified approach, we reasoned that both the mean methylation level and the mean *cov* of a feature (100kb bin or gene) could impact *mc* rate dispersion. We grouped features that fall into a combined bin of mean and *cov*. We then normalized the dispersion within each *mean-cov* group. After dispersion normalization, we selected the top 3000 features based on normalized dispersion for clustering analysis.

#### Dimension reduction and combination of different mC types

For each selected feature, *mc* rates were scaled to unit variance and zero mean. PCA was then performed on the scaled *mc* rate matrix. The number of significant PCs was selected by inspecting the variance ratio of each PC using the elbow method. The CH and CG PCs were then concatenated together for further analysis in clustering and manifold learning.

### Consensus clustering

#### Consensus clustering on concatenated PCs

We used a consensus clustering approach based on multiple Leiden-clustering^73^ over K-Nearest Neighbor (KNN) graph to account for the randomness of the Leiden clustering algorithms. After selecting dominant PCs from PCA in both mCH and mCG matrix, we concatenated the PCs together to construct a KNN graph using *scanpy*.*pp*.*neighbors* with euclidean distance. Given fixed resolution parameters, we repeated the Leiden clustering 300 times on the KNN graph with different random starts and combined these cluster assignments as a new feature matrix, where each single Leiden result is a feature. We then used the outlier-aware DBSCAN algorithm from the scikit-learn package to perform consensus clustering over the Leiden feature matrix using the hamming distance. Different epsilon parameters of DBSCAN are traversed to generate consensus cluster versions with the number of clusters that range from minimum to the maximum number of clusters observed in the multiple Leiden runs. Each version contained a few outliers that usually fall into three categories: 1. Cells located between two clusters that had gradient differences instead of clear borders, e.g., border of IT layers; 2. Cells with a low number of reads that potentially lack information in essential features to determine the specific cluster. 3. Cells with a high number of reads that were potential doublets. The amount of type 1 and 2 outliers depends on the resolution parameter and is discussed in the choice of the resolution parameter section. The type 3 outliers were very rare after cell filtering. The supervised model evaluation below then determined the final consensus cluster version.

#### Supervised model evaluation on the clustering assignment

For each consensus clustering version, we performed a Recursive Feature Elimination with Cross-Validation (RFECV)^74^ process from the scikit-learn package to evaluate clustering reproducibility. We first removed the outliers from this process, and then we held out 10% of the cells as the final testing dataset. For the remaining 90% of the cells, we used ten-fold cross-validation to train a multiclass prediction model using the input PCs as features and *sklearn*.*metrics*.*balanced_accuracy_score*^*75*^ as an evaluation score. The multiclass prediction model is based on *BalancedRandomForestClassifier* from the *imblearn* package that accounts for imbalanced classification problems^76^. After training, we used the 10% testing dataset to test the model performance using the *balanced_accuracy_score* score. We kept the best model and corresponding clustering assignments as the final clustering version. Finally, we used this prediction model to predict outliers’ cluster assignments, we rescued the outlier with prediction probability > 0.3, otherwise labeling them as outliers.

#### Manifold learning for visualization

In each round of clustering analysis, the t-SNE^77,78^ and UMAP^19^ embedding were run on the PC matrix the same as the clustering input using the implementation from the scanpy^72^ package. The coordinates from both algorithms were in Supplementary Table 5.

#### Choice of resolution parameter

Choosing the resolution parameter of the Leiden algorithm is critical for determining the final number of clusters. We selected the resolution parameter by three criteria: 1. The portion of outliers < 0.05 in the final consensus clustering version. 2. The ultimate prediction model accuracy > 0.9. 3. The average cell per cluster 30, which controls the cluster size to reach the minimum coverage required for further epigenome analysis such as DMR calls. All three criteria prevented the over-splitting of the clusters; thus, we selected the maximum resolution parameter under meeting the criteria using a grid search.

### Pairwise Differential Methylated Gene (DMG) identification

We used a pairwise strategy to calculate DMGs for each pair of clusters within the same round of analysis. We used the gene body ± 2kb regions of all the protein-coding and long non-coding RNA genes with evidence level 1 or 2 from the mouse GENCODE vm22. We used the per cluster normalized mCH rate (same as the “Computation and normalization of the methylation rate” in the clustering step above) to calculate markers between all neuronal clusters. We compared non-neuron clusters separately using the normalized mCG rate. For each pairwise comparison, we used the Wilcoxon test to select genes that have a significant decrease (hypo-methylation). Marker gene was chosen based on adjusted P-value < 1e-3 with multitest correction using Benjamini-Hochberg procedure, delta normalized methylation level change < −0.5 (hypo-methylation), AUROC > 0.8. We required each cluster to have 5 DMGs compared to any other cluster. Otherwise, the smallest cluster that did not meet this criterion was merged to the closest cluster based on cluster centroids euclidean distance in the PC matrix that was used for clustering. Then the marker identification process was repeated until all clusters found enough marker genes.

### Three-level of iterative clustering analysis

Based on the consensus clustering steps described above, we used an iterative approach to cluster the data into three levels of categories. In the first level termed CellClass, clustering analysis is done using all cells, and then manually merged into three canonical classes: excitatory neurons, inhibitory neurons, and non-neurons based on marker genes. Then within each CellClass, we performed all the preprocessing and clustering steps again to get clusters for the MajorType level using the same stop criteria. And within each MajorType, we obtained clusters for the SubType level. All clusters’ annotations and relationships are in Supplementary Table 4.

### Subtype phylogeny tree

To build the phylogeny tree of subtypes, we selected the top 50 genes that show the most number of significant changes for each subtypes’ pairwise comparisons. We then used the union of these genes from all subtypes and obtained a total of 2503 unique genes. We calculated the median mCH rate level of these genes in each subtype and applied bootstrap resampling based hierarchical clustering with average linkage and the correlation metric using the R package pvclust^79^ (v.2.2).

### Impact Score and Total Impact Score

We defined the Impact Score (IS) as a way to summarize pairwise comparisons for two subtype groups, where one group A contains *M* clusters, the other group B contains *N* clusters. For each gene or motif, the number of total related pairwise comparisons is *M* × *N*, the number of significant comparisons with desired change (hypo-methylation for gene or enrichment for motif) in group A is a, and in group B is *b*. The IS is then calculated as 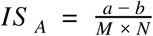 and 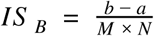 for the two directions. For either group, IS ranges from −1 to 1 and 0 means no impact, 1 means full impact, −1 means full impact in the other group.

We explored two scenarios of using the IS to describe cluster characteristics (Fig. 3e). The first scenario is considering each pair of branches in the subtype phylogeny tree as group A and group B. Thus the IS can quantify and rank genes or motifs to the upper nodes based on the leaves’ pairwise comparisons (Fig. 3i-h). The second scenario was a summarization of the total impact for specific genes or motifs regarding the phylogeny tree based on the calculation in the first scenario. In a subtype phylogeny tree with *n* subtypes, the total non-singleton node was *n-1*, and each node *i* had a height *h*_*i*_ and associated *lS*_*A*_ for one of the branches (*lS*_*B*_ *= - lS*_*A*_). The node height weighted total IS was then calculated by:

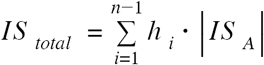

The larger total IS indicated a gene or motif show more cell-type-phylogeny related significant changes. The height weight intended to focus on the higher branches (major cell type separations), but the total IS can also be calculated in a sub-tree or any combination of interests, to rank gene and motifs most related to that combination (See Figure 5 related methods about calculating layer and region total IS from the same tree).

### Integration with snATAC-seq data

A portion of exact same brain tissue sample used in this study for methylome profiling was also processed with snATAC-seq in a parallel study about chromatin accessibility (Li et al., Companion Manuscript # 11). The final high-quality snATAC-seq cells were assigned to 160 chromatin accessibility clusters (a-clusters). The snATAC specific data analysis steps are described in Li et al. Here, we performed cross-modality data integration and label-transferring to assign the 160 a-clusters to the 161 methylome subtypes in the following steps:

1. We manually grouped both modalities into five integration groups (e.g., all IT neurons as a group) and only performed the integration of cells within the same group to decrease computation time. These groups were distinct in the clustering steps of both modalities and can be matched with great confidence using known marker genes. Step 2-6 were repeated for each group, see Extended Data Fig. 5 for the group design.
2. We used a similar approach as described above to identify pairwise differential accessible genes (DAG) between all pairs of snATAC-seq clusters. The cutoff for DAG is adjusted P-value < 1e-3, fold change > 2, AUROC > 0.8.
3. We then gather DMGs from related subtypes’ comparisons in the same group. Both DAGs and DMGs were filtered by recurring in > 5 pairwise comparisons. The intersection of the remaining genes was used as the feature set of integration.
4. After identifying DAGs using cell level snATAC-seq data, we merged the snATAC-seq cells into pseudo-cells to increase snATAC-seq data coverage. Within each a-cluster, we did a K-means clustering (K = # of cells in that cluster / 50) on the same PCs that were used in snATAC-seq clustering. We discarded (about 5% of the cells) small K-means clusters with < 10 cells and merged each remaining K-means cluster into a pseudo-cell. Each pseudo-cell had about 50 times more fragments than a single cell.
5. We then used the MNN based Scanorama^39^ method with default parameters to integrate the snmC-seq cells and snATAC-seq pseudo-cells using genes from 3. After Scanorama integration, we did co-clustering on the integrated PC matrix using the clustering approaches described above.
6. We used the intermediate clustering assignment from 5 to calculate the overlap score (see the section below) between the original methylome subtypes and the a-clusters. We used the overlap score > 0.3 to assign snATAC-seq clusters to each methylome subtype. For those subtypes that have no match under this threshold, we assign the top a-cluster ranked by the overlap score.

### Overlap Score (OS)

We used the overlap score to match a-cluster and methylome subtypes together. The overlap score range from 0 to 1 was defined as the sum of the minimum proportion of samples in each cluster that overlapped within each co-cluster^26^. A higher score between one methylome subtype and one a-cluster indicates they consistently co-clustered within one or more co-clusters. Besides matching clusters in integration analysis, the OS was also used in two other cases: 1. To quantify replicates and region overlaps over methylome subtypes (Extended Data Fig. 2e-g); 2. To quantify the overlap of each L5-ET subtype overlapping with “soma location” and “projection target” labels from epi-retro-seq cells (Extended Data Fig. 4k) through integration with the epi-retro-seq dataset.

## STAGE III. CELL-TYPE-SPECIFIC REGULATORY ELEMENTS

### Differentially Methylated Region (DMR) analysis

After clustering analysis, we used the subtype cluster assignments to merge single-cell ALLC files into the pseudo-bulk level and then used methylpy^70^ *DMRfind* function to calculate mCG DMRs across all clusters. The base calls of each pair of CpG sites were added before analysis. In brief, the methylpy function used a permutation-based root-mean-square test of goodness-of-fit to identify differentially methylated sites simultaneously across all samples (subtypes in this case), and then merge the DMS within 250bp into DMR. We further excluded DMS calls that have low absolute mCG rate difference by using a robust-mean-based approach. For each DMR that was merged from the DMS, we order all the samples by their mCG rate and calculate the robust mean *m* using the samples between 25th and 75th percentiles. We then reassign hypo-DMR and hyper-DMR to each sample when a region met two criteria: 1) the sample mCG rate of this DMR is lower than (*m* - 0.3) for hypo-DMR or (*m* + 0.3) for hyper-DMR, and 2) the DMR is originally a significant hypo- or hyper-DMR in that sample judged by methylpy. DMRs without any hypo- or hyper-DMR assignment were excluded from further analyses.

### Enhancer prediction using DNA methylation and chromatin accessibility

We performed enhancer prediction using the REPTILE^51^ algorithm. The REPTILE is a random-forest-based supervised method that incorporates different sources of epigenomic profiles with base-level DNA methylation data to learn and then distinguish the epigenomic signatures of enhancers and genomic background. We trained the model in a similar way as in the previous studies^6,51^ using CG methylation, chromatin accessibility of each subtype, and mouse embryonic stem cells (mESC). The model was first trained on mESC data and then predicted a quantitative score we termed enhancer score for each subtype’s DMRs. The positives were 2kb regions centered at the summits of top 5,000 EP300 peaks in mESCs. Negatives include randomly chosen 5,000 promoters and 30,000 2kb genomic bins. The bins have no overlap with any positive region or gene promoter^6^.

Methylation and chromatin accessibility profiles in bigwig format for mESC were from the GEO database (GSM723018). The mCG rate bigwig file was generated from subtype-merged ALLC files using the ALLCools package (https://github.com/lhqing/ALLCools). For chromatin accessibility of each subtype, we merged all fragments from snATAC-seq cells that were assigned to this subtype in the integration analysis and used “*deeptools bamcoverage*” to generate CPM normalized bigwig files. All bigwig file bin sizes were 50bp.

### Motif enrichment analysis

We used 719 motif PWMs from the JASPAR 2020 CORE vertebrates database^43^, where each motif was able to assign corresponding mouse TF genes. The specific DMR sets used in each motif enrichment analysis are described in figure specific methods below. For each set of DMRs, we standardized the region length to the center ± 250bp and used the FIMO tool from the MEME suite^80^ to scan the motifs in each enhancer with the log-odds score p-value < 10^−6^ as the threshold. To calculate motif enrichment, we use the adult non-neuronal mouse tissue DMRs^65^ as background regions unless expressly noted. We subtracted enhancers in the region set from the background, and then scanned the motifs in background regions using the same approach. We then used Fisher’s exact test to find motifs enriched in the region set, and the Benjamini-Hochberg procedure to correct multiple tests. We used the TFClass^81^ classification to group TFs with similar motifs.

### DMR-DMG partial correlation

To calculate DMR-DMG partial correlation, we used the mCG rate of DMRs and the mCH rate of DMGs in each neuronal subtypes. We first used linear regression to regress out variance due to global methylation difference (using scanpy.pp.regress_out function), then use the residual matrix to calculate Pearson correlation between DMR and DMG pairs where the DMR center is within 1Mb the DMG’s TSSs. To generate the null distribution, we shuffled the subtype orders in both matrices and recalculated all pairs 100 times.

### Identification of loops and differential loops from sn-m3C-seq data

After merging the chromatin contacts from cells belonging to the same type, we generated a .hic file of the cell-type with *Juicer tools pre. HlCCUPS*^*82*^ was used to identify loops in each cell-type. The loops from eight major cell-types were concatenated and deduplicated, and used as the total samples for differential loop calling. A loop-by-cell matrix was generated, where each element represents the number of contacts supporting each loop in each cell. The matrix was used as input of EdgeR to identify differential interactions with ANOVA tests. Loops with FDR < 1e-5 and min-max fold-change > 2 were used as differential loops.

## FIGURE-SPECIFIC METHODS

**Figure 1 related**

### 3D model of dissection regions (Fig. 1d-g)

We created in-silico dissection regions based on the Allen CCFv3^69^ 3D model using blender 2.8 that precisely follows our dissection plan. To ease visualization of all different regions, we modified the layout and removed some of the symmetric structures, but all the actual dissections were applied symmetrically to both hemispheres.

### Calculating the genome feature detected ratio (Fig. 1i)

The detected ratio of chromosome 100kb bins and gene bodies is calculated as the percentage of bins with > 20 total cytosine coverage. Non-overlapping chromosome 100kb bins generated by “bedtools makewindows -w 100000”; gene body definition from the GENCODE vm22 GTF file.

**Figure 2 related**

### Integration with epi-retro-seq L5-ET cells (Fig. 2i-l, Extended Data Fig. 4h-j)

Epi-retro-seq is an snmC-seq2 based method that combines retrograde AAV labeling (Companion Manuscript # 11)^36^. The L5-ET cells’ non-overlapping chromosome 100kb bin matrix gathered by the epi-retro-seq dataset was concatenated with all the L5-ET cells from this study to do co-clustering and embedding as described in “STAGE II” above. We then calculated the OS between subtypes in this study and the “soma location” or “projection target” labels of epi-retro-seq cells. The first OS helped quantify how consistent the spatial location is between the two studies, the second OS allowed us to impute the projection targets of subtypes in this study.

**Figure 3 related**

### Pairwise DMR and motif enrichment analysis (Fig. 3c, h)

The total subtype DMRs were identified as described in “STAGE III” via comparing all subtypes. We then assigned DMRs to each subtype pair if the DMRs were: 1) significantly hypomethylated in only one of the subtypes; and 2) the mCG rate difference between the two subtypes > 0.4. Each subtype pair was associated with two exclusive sets of pairwise DMRs. We carried out motif enrichment analysis as described in “STAGE III” on each DMR set using the other set as background. Motifs enriched in either direction were then used to calculate the impact score and were associated with upper nodes of the phylogeny.

**Figure 4 related**

### Overlapping eDMR with genome regions (Fig. 4b)

The cluster-specific snATAC-seq peaks were identified in Li et al. (Companion Manuscript # 11). We used “bedtools merge” to aggregate the total non-overlap peak regions, and “bedtools intersect” to calculate the overlap between peaks and eDMRs. The developing forebrain and other tissue feDMR were identified in He et al.^6^ using methylC-seq^83^ for bulk whole-genome bisulfite sequencing (WGBS-seq). All of the genome features used in Fig. 4b were defined as in He et al, except using an updated mm10 CGI region and RepeatMaster transposable elements lists (UCSC table browser downloaded on 09/10/2019, and the GENCODE vm22 gene annotation).

### Assembly epigenome card (Fig. 4f) of the gene-enhancer landscape

The eDMRs for each gene selected by eDMR-Gene correlation > 0.3. Sections of the heatmaps in Fig. 4f were gathered by: 1) mCG rate of each eDMR in 161 subtypes from this study, 2) snATAC subtype-level FPKM of each eDMR in the same subtype orders. The subtype snATAC profiles were merged from integration results as described in “STAGE II”, 3) mCG rate of each eDMR in forebrain tissue during ten developing time points from E10.5 to P0, data from He et al.^6^, 4) H3K27ac FPKM of each eDMR in 7 developing time points from E11.5 to P0, data from Gorkin et al.^68^, 5) H3K27ac FPKM of each eDMR in P56 frontal brain tissue, data from Lister et al.^2^ and 6) eDMR is overlapped with forebrain feDMR using “bedtools intersect”.

### Embedding of cells with chromosome interactions (Fig. 4i)

scHiCluster^84^ was used to generate the t-SNE embedding of the sn-m3C-seq cells. Specifically, a contact matrix at 1Mb resolution was generated for each chromosome of each cell. Then the matrices were smoothed by linear convolution with pad = 1 and random walk with restart probability = 0.5. The top 20th percentile of strongest interactions on the smoothed map was extracted, binarized, and used for PCA. The first 20 PCs were used for t-SNE.

**Figure 5 related**

### IT layer-dissection-region group DMG and DMR analysis (Fig. 5a-f)

In order to collect enough cells for dissection region analysis, we only used the major types (which corresponds to L2/3, L4, L5, and L6) of IT neurons. We grouped cells into layer-dissection-region groups and kept groups with > 50 cells in further analysis (Extended Data Fig. 8b). We performed pairwise DMG, DMR, and motif enrichment analysis the same as the subtype analysis in Fig. 3, but using the layer-dissection-region group labels. We then built a spatial phylogeny for these groups and calculated impact scores based on it. To rank layer-related or dissection-region-related genes and motifs separately, we used two sets of the branches (Extended Data Fig. 8a, upper set for layers, lower set for regions) in the phylogeny and calculated two total impact scores using equations in above.

### DG cell group and gradient DMR analysis (Fig. 5h)

DG cells were grouped into four even-sized groups according to cells’ global mCH rates, and cutoff thresholds were 0.45%, 0.55%, and 0.69%. We then randomly chose 400 cells from each group to call gradient-DMRs using methods described in “STAGE II”. To ensure the DMRs identified between intra-DG groups were not due to stochasticity, we also randomly sampled 15 groups of 400 cells from all DG cells regardless of their global mCH and called DMR among them as control-DMRs (2,003 using the same filtering condition). Only 0.04% of gradient-DMRs overlapped with the control-DMRs and thus were removed from further analysis. Pearson correlation coefficients (p’s) of mCG rates of each gradient-DMR were calculated against a linear sequence [1, 2, 3, 4] to quantify the gradient trend. DMRs with p<-0.75 or p>0.75 were considered to be significantly correlated. Weakly correlated DMRs (10%) were not included in further analysis.

### DMR/DMS enriched genes (Fig. 5i-j)

To investigate the correlated DMR/DMSs enrichment in specific gene bodies, we compare the number of DMS and cytosine inside the gene body with the number of DMS and cytosine in the ± 1Mb regions through Fisher’s exact test. We chose genes passing both criteria: 1) adjusted P-value < 0.01 with multitest correction using Benjamini-Hochberg procedure, and 2) overlap with > 20 DMS. Gene ontology analysis of DMR/DMS enriched genes was carried out using GOATOOLS^85^ all protein coding genes with gene body length > 5kb were used as background to prevent gene length bias.

### Compartment strength analysis

We Z-score normalized the total chromosome contacts in each 1 Mb bin of DG contact matrix, and the bins with normalized coverage between −1 and 2 were kept for the analysis. After filtering, the PC1 of genome-wide KR normalized contact matrix was used as the compartment score. The score was divided into 50 categories with equal sizes from low to high, and bins were assigned into the categories. The intra-chromosomal observation/expectation (ove) matrices of each group were used to quantify the compartment strength. We computed the average ove values within each pair of categories to generate the 50×50 saddle matrices. The compartment strength was computed with the average of upper left and lower right 10×10 matrices divided by the average of upper right and lower left 10×10 matrices^86^.

### Domain analysis

We identified 4,580 contact domains at 10kb resolution in DG using *Arrowhead*^*82*^. For bin, the insulation score is computed by

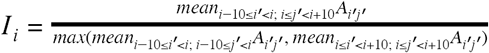

where *A* is the ove of KR normalized matrices. For each group, insulation scores of domain boundaries and 100kb flanking regions were computed and averaged across all boundaries.

**Figure 6 related**

#### Prediction model description

To reduce the computing complexity, we applied principal component analysis (PCA) on the dataset of 100kb-bin-mCH features to obtain the first 3,000 principal components (PCs), which retains ∼ 61% variance of the original data. These 3,000 PCs were then used to train and test the predicting model. We used an artificial neural network with two hidden layers to predict cell subtypes and their dissection regions simultaneously (Fig. 6a). The input layer contains 3,000 nodes, followed by a shared layer with 1,000 nodes. The shared layer is further connected simultaneously to two branch hidden layers of the subtype of the dissection region, each containing 200 nodes. Branch hidden layers are followed by the corresponding one-hot encoding output layers. We used 5-fold cross-validation to access the model performance. During the training, we applied the dropout technique^87^ with a dropout rate *p*=0.5 on each hidden layer to prevent overfitting. Adam optimization^88^ was used to train the network with a cross-entropy loss function. The training epoch number and batch size are 10 and 100, respectively. The training and testing processes were conducted via TensorFlow 2.0^89^.

#### Model performance

For each single cell input, the two output layers generate two probabilistic vectors as the prediction results for cell subtypes and dissection regions, respectively. The subtype and dissection region label with highest probabilities were used as the prediction results for each cell to calculate accuracy. When calculating the cell dissection region accuracy (Fig. 6c), we defined two kinds of accuracy with different stringency: 1) the exact accuracy using the predicted label, and 2) the fuzzy accuracy using predicted labels or its potential overlap neighbors. The potential overlap neighbors curated based on Allen CCF (Extended Data Fig. 9a, Supplementary Table 2) stood for adjacent regions of a particular dissection region. The exact accuracy of the ANN model is 69%, and the fuzzy accuracy is 89%. To evaluate how much of the dissection region accuracy was improved via ANN, we calculated fuzzy accuracy just based on naive guesses in each subtype based on the dissection region composition (gray dots in Extended Data Fig. 9b).

#### Biological feature importance for dissection region prediction

To assess what DNA regions store information of cell spatial origins that is distinguishable using our model, we evaluated the PC feature importance by examining how permutation of each PC feature across cells affects prediction accuracy. We did five permutations for each feature and used the average accuracy decreasing as PC feature importance. For a given cell type, we examined genes contained in the 100kb bins with the top 1% PCA factor loadings for the most important PC feature.

## Data availability

Single cell raw and processed data included in this study were deposited to NCBI GEO/SRA with accession number GSE132489 and to the NeMO archive: https://portal.nemoarchive.org/. Cluster merged methylome profiles can be visualized at http://neomorph.salk.edu/mouse_brain.php.

## Code availability

The mapping pipeline for snmC-seq2 data: https://cemba-data.readthedocs.io/en/latest/; The ALLCools package for post-mapping analysis and snmC-seq2 related data structure: https://github.com/lhqing/ALLCools; The jupyter notebooks for reproducing specific analysis: https://github.com/lhqing/mouse_brain_2020.

## Supplementary Tables

Supplementary Table 1. Glossary table for all the abbreviations.

Supplementary Table 2. Metadata of the 45 brain dissection regions.

Supplementary Table 3. Summary of cell numbers.

Supplementary Table 4. Cell type names and annotations.

Supplementary Table 5. Cell metadata and manifold learning coordinates.

Supplementary Table 6. Cell type by dissection region cell counts.

Supplementary Table 7. snATAC-seq clusters matching to subtypes.

---

**Extended Data Fig. 1.**
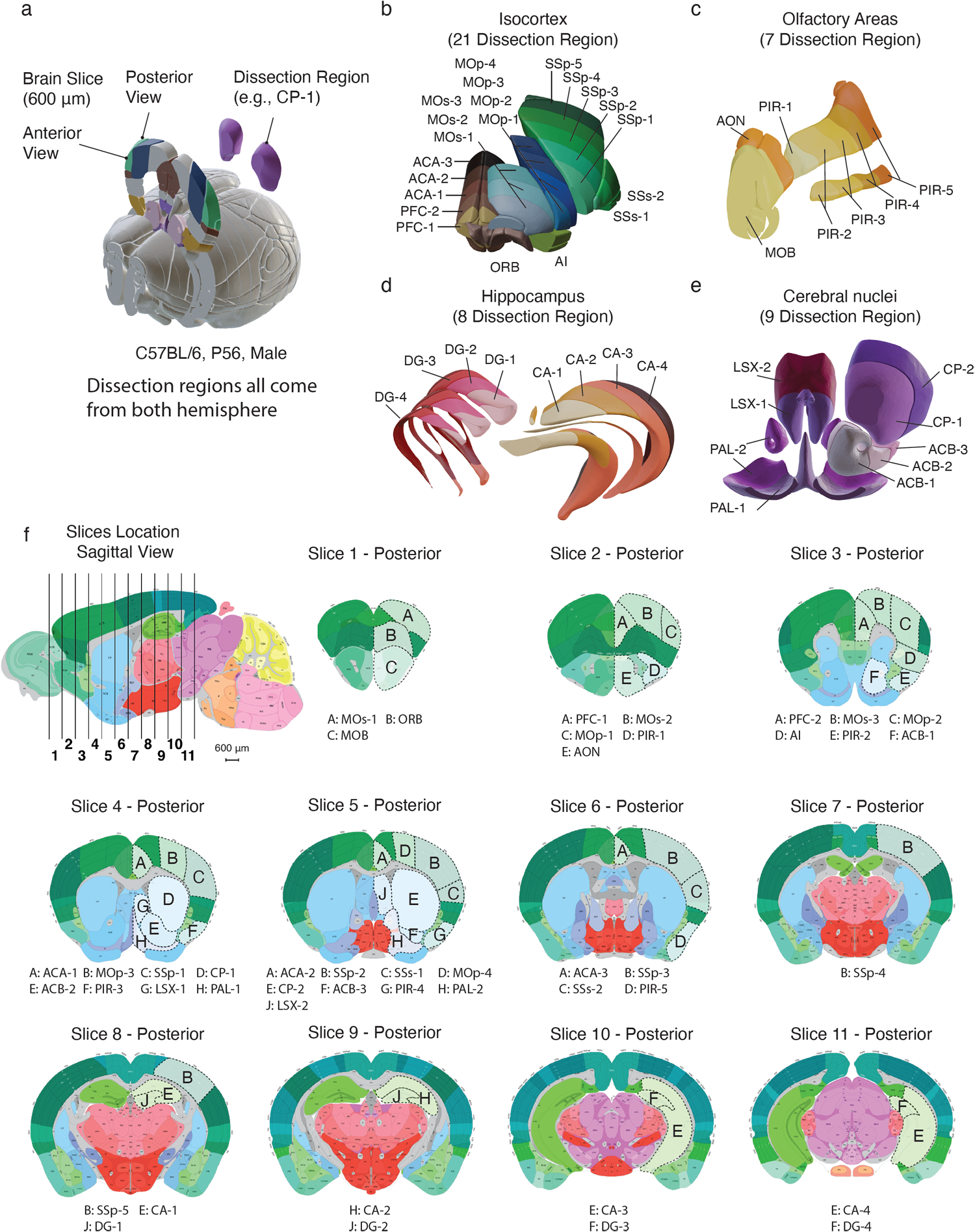
Brain dissection regions. **a**, 3D schematic of brain dissection steps. Each P56, male, C57BL/6 mouse brain is dissected into 600-micron slices. we then dissect brain regions from both hemispheres within a specific slice. **b-e**, 3D adult P56 mouse brain schematic adapted from Allen CCFv3 to display the four major brain regions and 45 dissection regions. Each color and the corresponding label represents a dissection region. **f**, 2D adult P56 mouse brain atlas adapted from Allen Mouse Brain Reference Atlas, the first sagittal image showing the location of each coronal slice, followed by 11 posterior view images of all coronal slices, the same 45 dissection regions are labeled on the corresponding slice. All coronal images follow the same scale as the sagittal image. The posterior view of each slice is the anterior view of the next slice.

**Extended Data Fig. 2.**
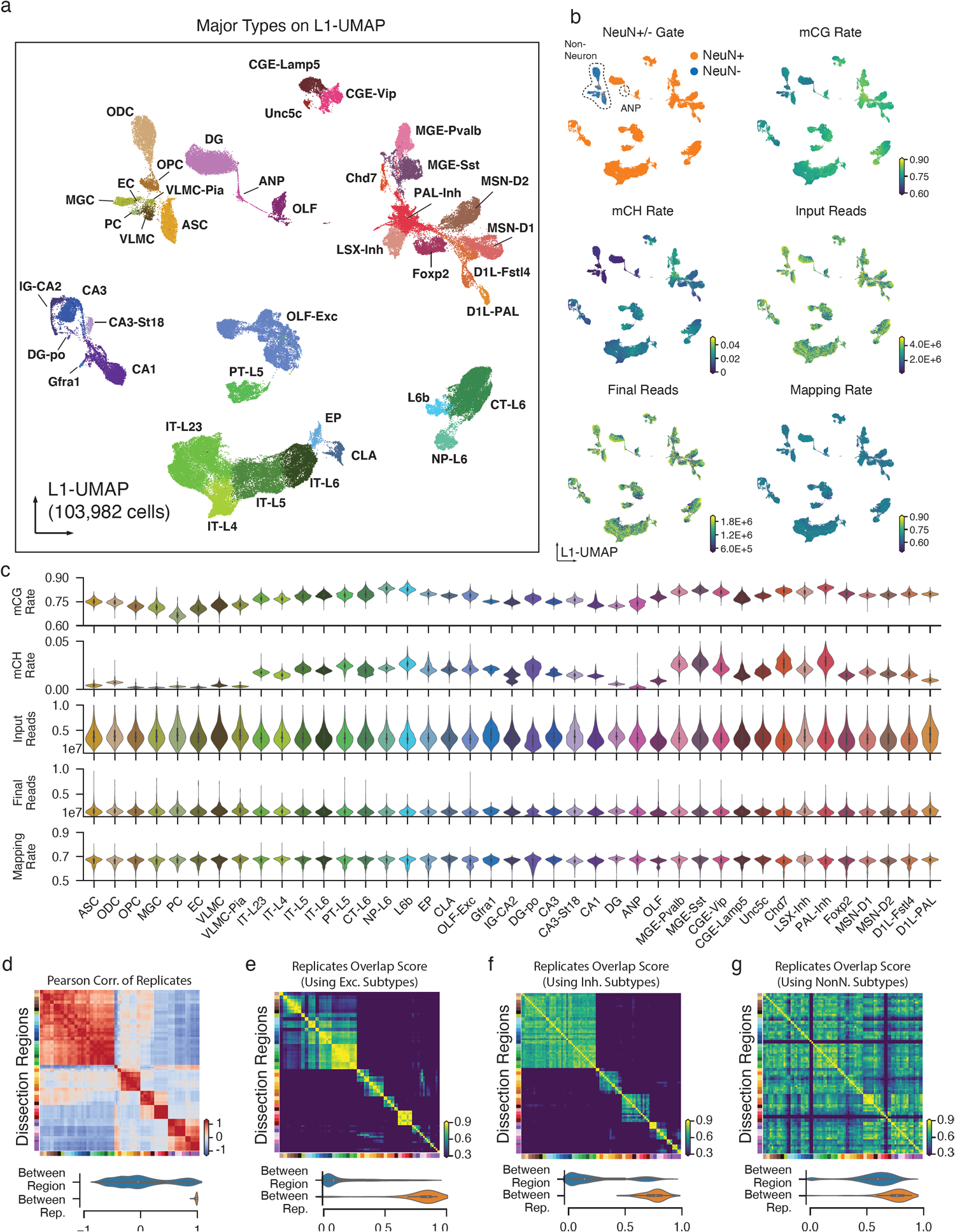
Major Type labeling and basic mapping metrics of snmC-seq2. **a**, L1-UMAP colored and labeled by major cell types. **b**, L1-UMAP colored by NeuN antibody FACS gates and other snmC-seq2 key read mapping metrics. **c**, Violin plots for all of the key metrics, group by major types. **d**, Heatmap of Pearson correlation between the average methylome profiles (mean mCH and mCG rate of all chromosome 100kb bins across all cells belong to a replicate sample) of the 92 replicates from 45 brain regions. The violin plot below summarizes the value between replicates within the same brain region, or between different brain regions. **e-g**, Pairwise overlap score (measuring co-clustering of two replicates) of excitatory subtypes (**e**), inhibitory subtypes (**f**), and non-neuronal subtypes (**g**). The violin plots summarize the subtype overlap score between replicates within the same brain region, or between different brain regions.

**Extended Data Fig. 3.**
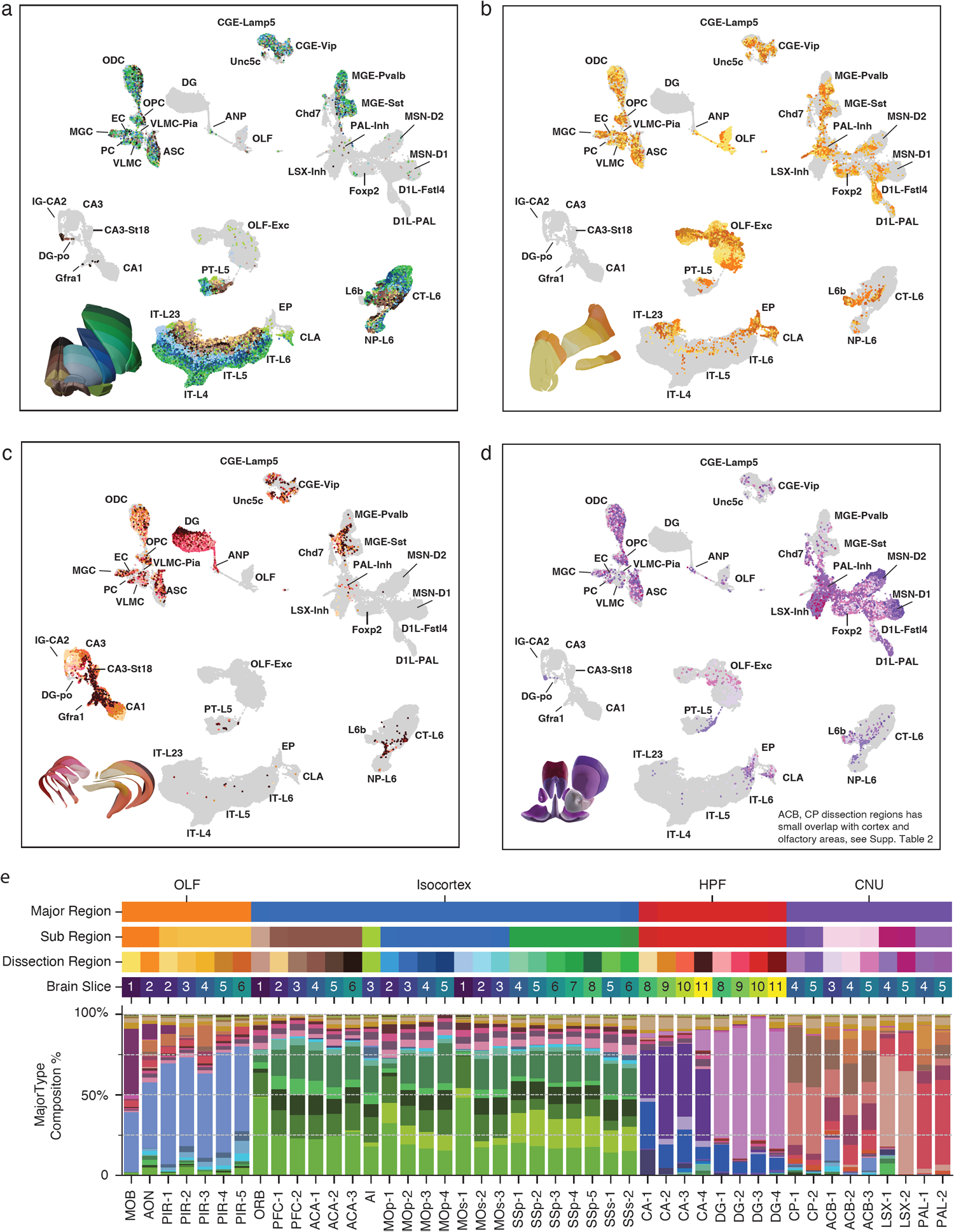
Cell-type composition of dissection regions. **a-d**, L1-UMAP labeled by major types and partially colored by dissection regions for cells from Isocortex (**a**), olfactory areas (**b**), hippocampus (**c**), and cerebral nucleus (**d**). Other cells are shown in gray as background. **e**, Similar compound bar plot as Fig. 1h, from top to bottom, showing the organization of dissection regions and the major type composition of each dissection region.

**Extended Data Fig. 4.**
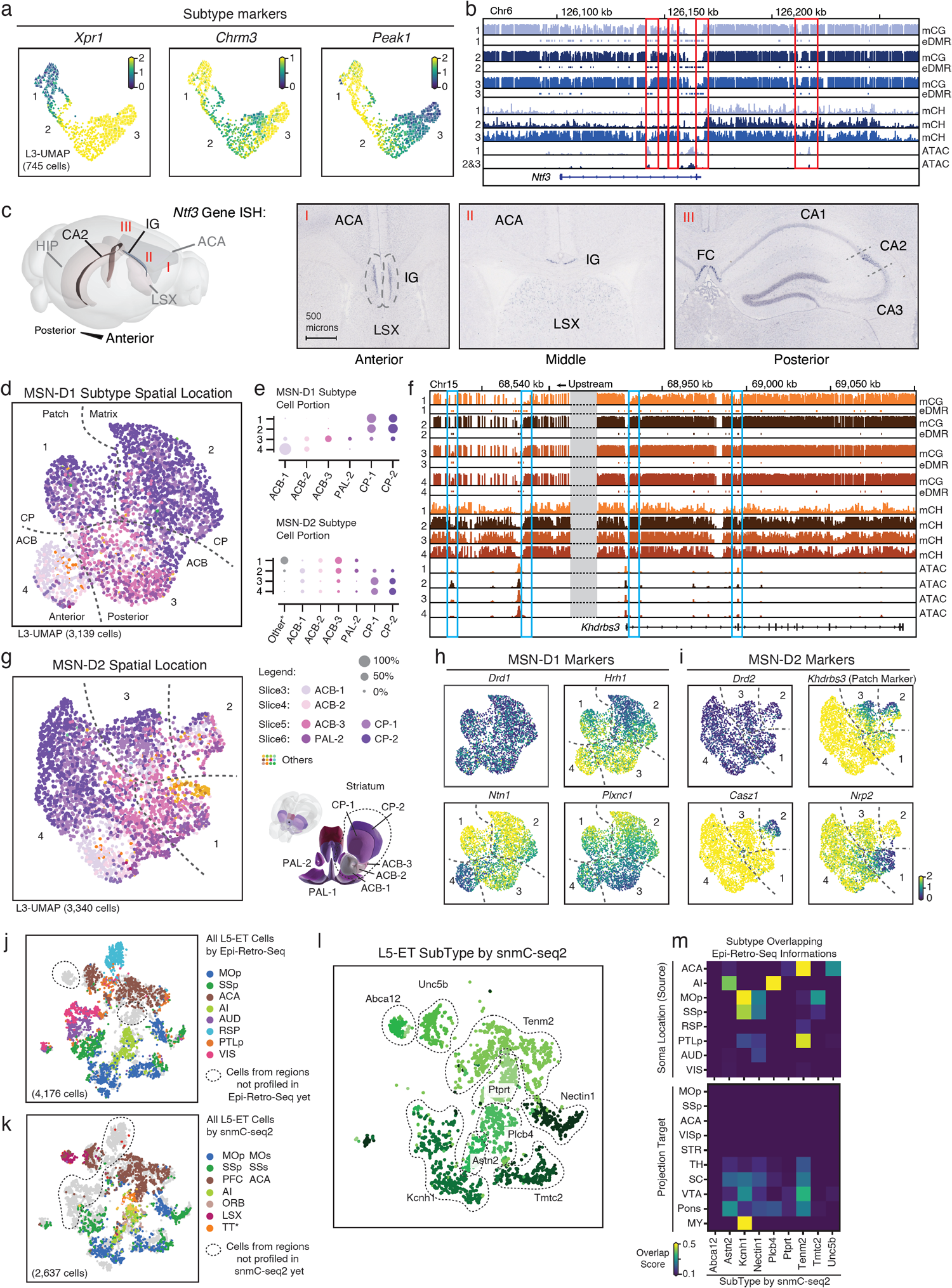
Supporting details of cellular and spatial diversity of neurons at the subtype level. **a**, mCH Rate of marker genes in IG-CA2 cells. **b**, Methylome and chromatin accessibility genome browser view of *Ntf3* genes and its upstream regions. ATAC and eDMR information are from Fig.4 analysis. **c**, Three different views of the ISH experiment from ABA, showing the *Ntf3* gene expressed in both IG and CA2. **d**, L3-UMAP from MSN-D1 cells colored by subregion. The numbers are subtypes marked by hypomethylation of unique genes: 1) *Khdrbs3*, 2) *Hrh1*, 3) *Plxnc1*, an 4) *Ntn1*. **e**, The dot plot shows region composition of each subtype of MSN-D1 and MSN-D2. **f**, Genome browser view of *Khdrbs3* genes similar to (**b**). **g**, MSN-D2 subtypes marked by hypomethylation of unique genes: 1) *Nrp2*, 2) *Casz1*, 3) *Col14a1*, and 4) *Slc24a2*. **h, i**, mCH Rate of MSN-D1 (**h**) and MSN-D2 (**i**) subtype marker genes. **j, k**, Same integration t-SNE as Fig. 2i-j colored by the dissection regions, but using all cells that have been profiled by Epi-Retro-Seq (**j**) or snmC-seq2 (**k**), cells from brain regions that have only been profiled via one of the methods are circled out. **l**, Same t-SNE as (**k**) colored subtypes. **m**, Overlap score matrix matching the subtypes to the “Soma Location (source)” and “Projection Target” information labels of Epi-Retro-Seq cells.

**Extended Data Fig. 5.**
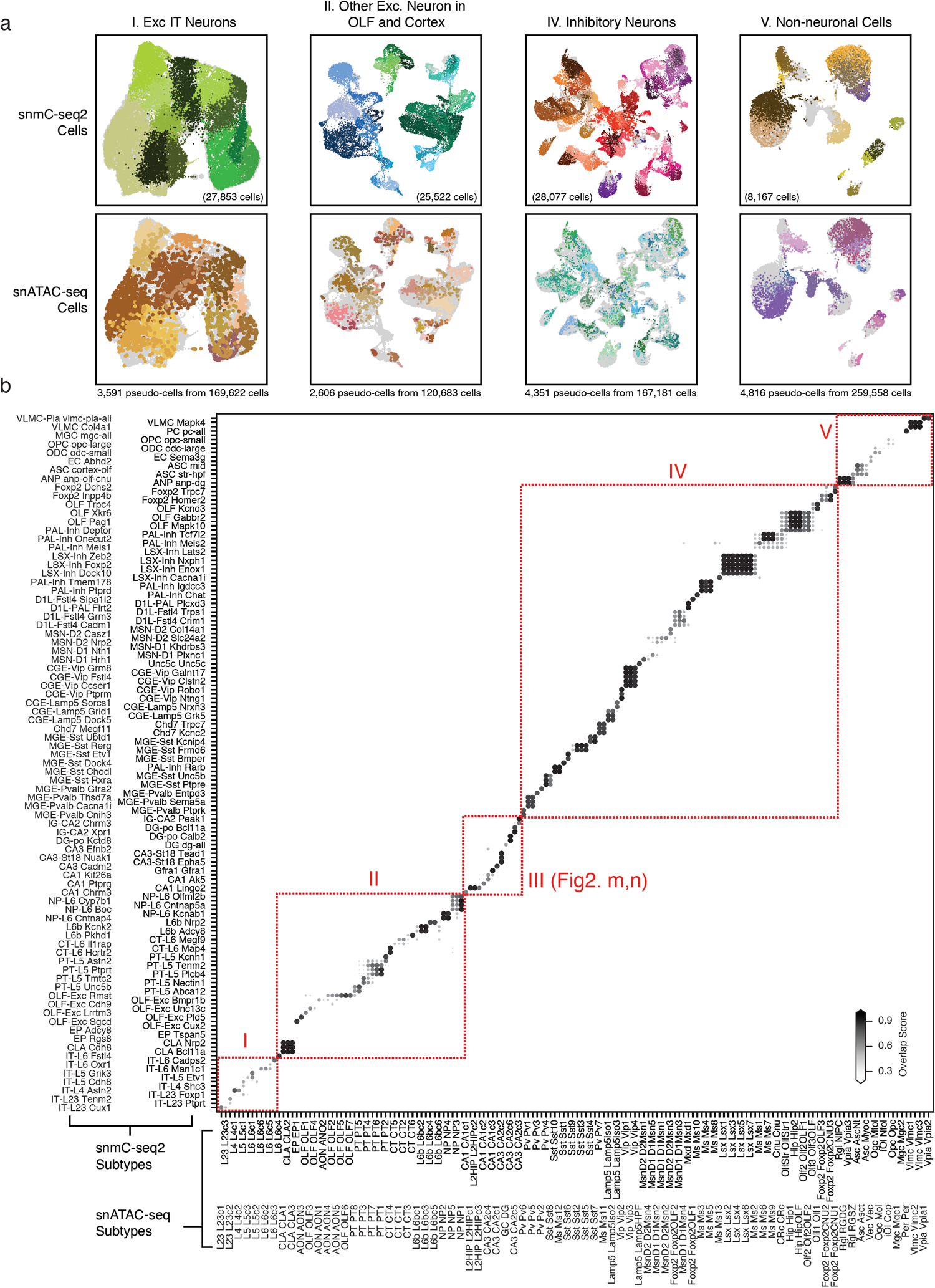
Integration with snATAC-seq. **a**, Integration UMAP for snmC-seq2 cells (top row) and snATAC-seq pseudo-cells (bottom row). Each panel is colored by subtypes from the corresponding study, the other dataset shown in gray in the background. snATAC-seq subtype color token from Li et al., (Companion Manuscript # 11). Integration is done in five separate groups, including four shown here and one shown in Fig. 2m-n. **b**, Overlap score matrix matching the 160 snATAC-seq subtypes to the 161 snmC-seq2 subtypes.

**Extended Data Fig. 6.**
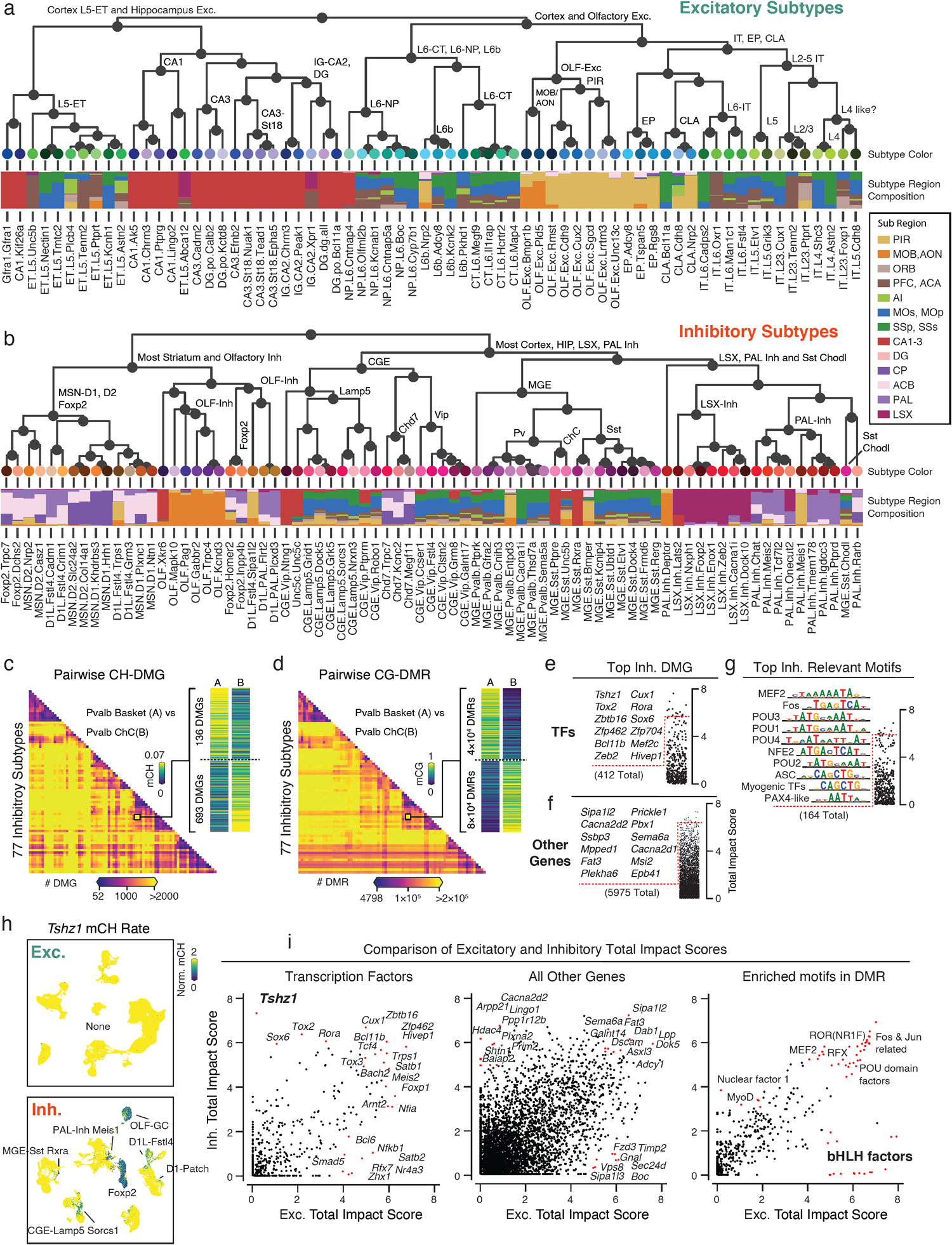
Subtype phylogeny with related gene and motifs. **a, b**, Subtype phylogeny of excitatory (**a**) and inhibitory (**b**) neurons. Leaf nodes colored by subtypes, and the barplot shows subregion composition. **c, d**, Counts heatmap of pairwise CH-DMG (**c**) and CG-DMR (**d**) between 77 inhibitory subtypes. **e-g**, Top TFs (**e**), other genes (**f**), and enriched motifs (**g**) rank by total impact score based on the inhibitory subtype phylogeny. **h**, An example gene Tshz1 only shows subtype diversity in inhibitory subtypes but not excitatory subtypes. **i**, Comparison of the total impact scores calculated from either excitatory subtype phylogeny (x-axis) or inhibitory subtype phylogeny (y-axis) for TFs, other genes, and enriched motifs.

**Extended Data Fig. 7.**
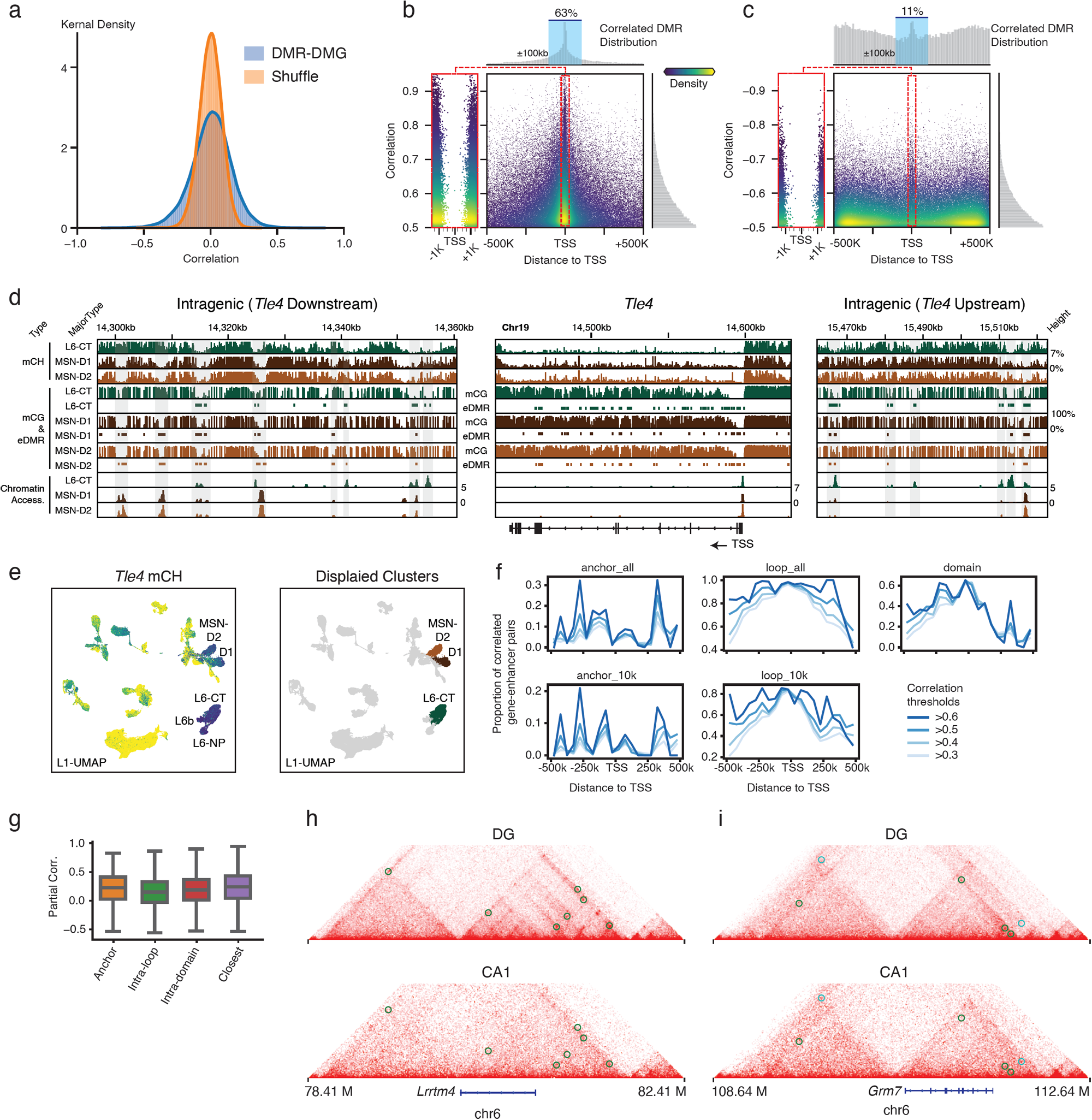
Gene-Enhancer landscape related. **a**, Distribution of actual DMR-DMG partial correlation compared to the shuffled null distribution. **b, c**, DMR-DMG correlation (y-axis) and the distance between DMR center and gene TSS (x-axis), each point is a DMR-DMG pair, color represents points kernel density. The positively (**b**) and negatively (**c**) correlated DMRs are shown separately, due to very different genome location distributions that are plotted on the top histograms. **d**, detailed view of surrounding eDMRs that are correlated with *Tle4* gene body mCH. Alternative eDMRs only appear in either L6-CT or MSN-D1/D2 can be seen on both upstream and downstream of the gene. **e**, *Tle4* gene body mCH and the major type label on L1-UMAP, paired with (**d**). **f**, Proportion of loop supported enhancer-gene pairs among those linked by correlation analyses surpassing different correlation thresholds at each specific distance. **g**, Partial correlation between mCG of enhancers and mCH of genes linked by different methods (n=4,171, 127,730, 28,203, 10,058). The elements of boxplots are defined as: center line, median; box limits, first and third quartiles; whiskers, 1.5x interquartile range. **h, i**, Interaction maps, mCH, mCG, ATAC, and differential loops tracks surrounding *Lrrtm4* (**h**) and *Grm7* (**i**). Circles on the interaction maps represent differential loops between DG and CA1, where green represents DG loops, and cyan represents CA1 loops.

**Extended Data Fig. 8.**
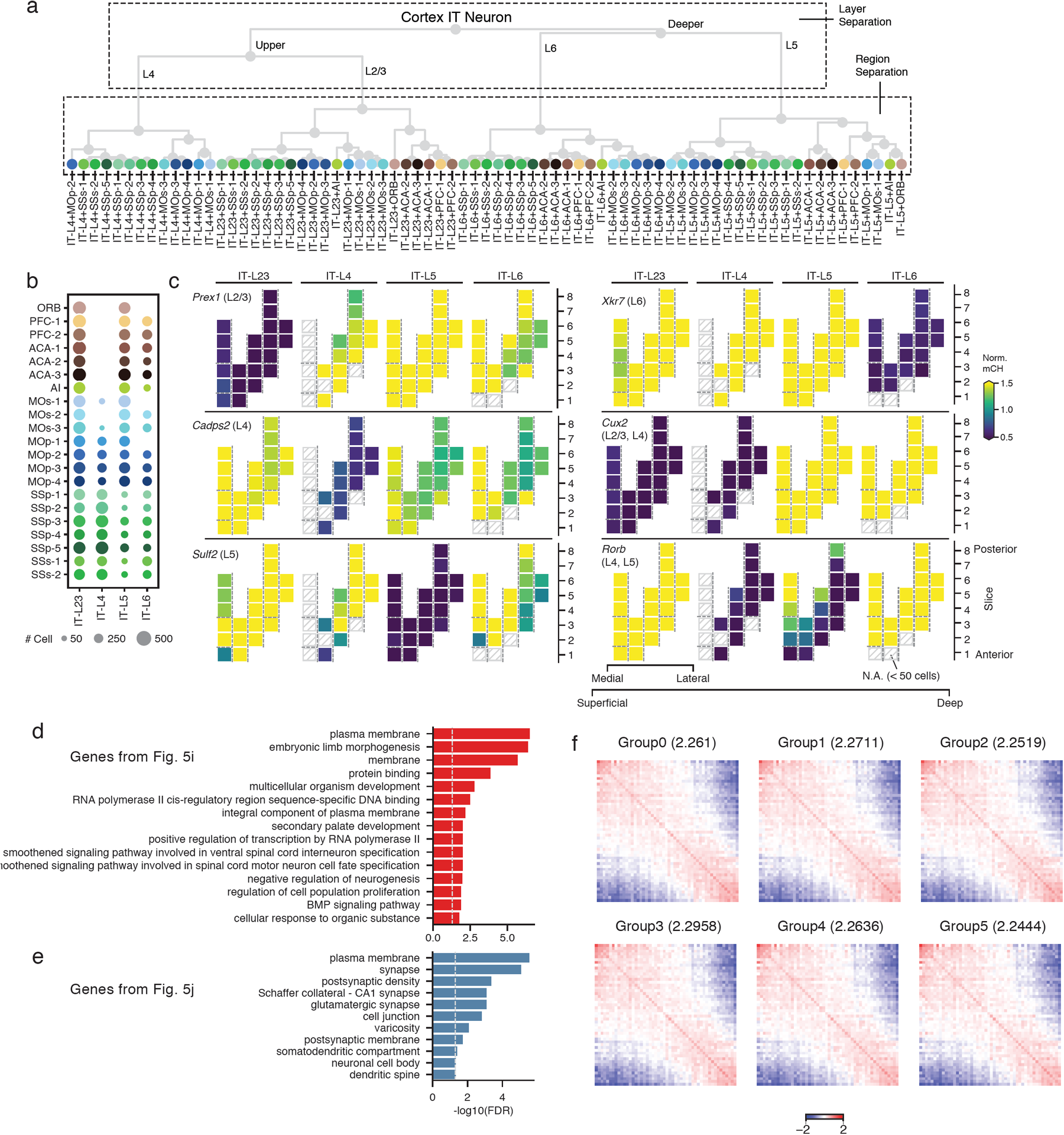
DNA methylation gradient related. **a**, Layer-dissection-region cell group phylogeny. **b**, Dot plot sized by the number of cells in each layer-dissection-region combination in excitatory IT neurons. Each group needs at least 50 cells to be included in the analysis. **c**, Representative marker genes for laminar layers separation. Using the same dissection region layout as Fig. 5b. **d, e**, Top enriched GO terms for positively (**d**) and negatively (**e**) correlated DMR enriched genes. **f**, Saddle plots of different groups of DG cells separated by global mCH. Values in the title represent the compartment strengths.

**Extended Data Fig. 9.**
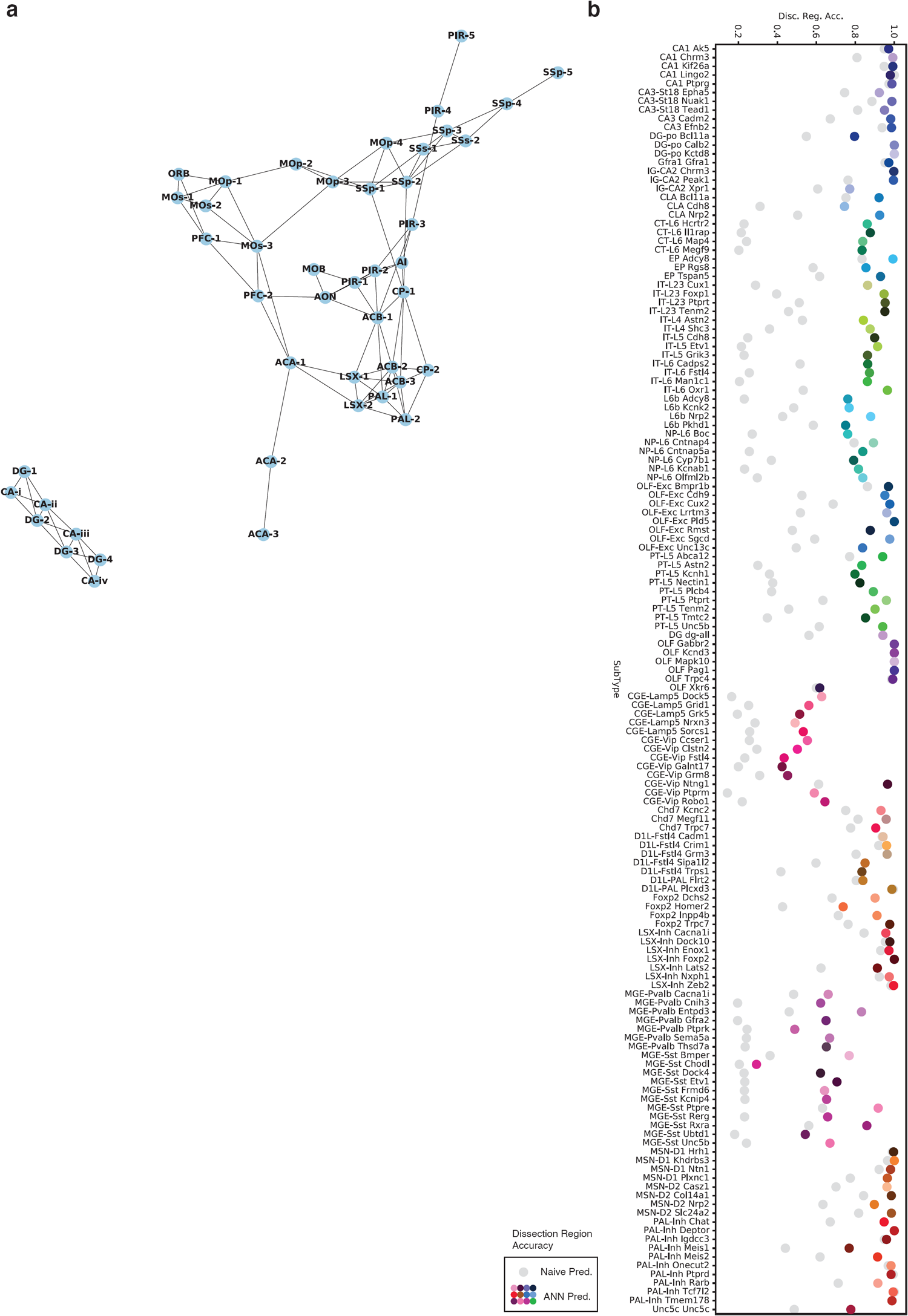
Evaluation of the predictive model. **a**, The neighbor relation among the potential overlapping dissection regions. The network is constructed based on information of the dissection scheme and the “Potential Overlap” column in Supplementary Table 2. **b**, Prediction accuracy of dissection region at cell subtype level. The colored points denote the prediction accuracy of the model, while the grey ones denote the random guess accuracy when cell subtypes and corresponding spatial distributions are given.

## Reference

1. Luo, C., Hajkova, P. & Ecker, J. R. Dynamic DNA methylation: In the right place at the right time. Science 361, 1336–1340 (2018).

2. Lister, R. et al. Global epigenomic reconfiguration during mammalian brain development. Science 341, 1237905 (2013).

3. Fagiolini, M., Jensen, C. L. & Champagne, F. A. Epigenetic influences on brain development and plasticity. Curr. Opin. Neurobiol. 19, 207–212 (2009).

4. Lavery, L. A. et al. Losing Dnmt3a dependent methylation in inhibitory neurons impairs neural function by a mechanism impacting Rett syndrome. Elife 9, (2020).

5. Li, J., Pinto-Duarte, A., Zander, M., Lai, C. Y. & Osteen, J. Polycomb-mediated repression compensates for loss of postnatal DNA methylation in excitatory neurons. bioRxiv (2019).

6. He, Y. et al. Spatiotemporal DNA Methylome Dynamics of the Developing Mammalian Fetus. bioRxiv 166744 (2017) doi:10.1101/166744.

7. Mo, A. et al. Epigenomic Signatures of Neuronal Diversity in the Mammalian Brain. Neuron 86, 1369–1384 (2015).

8. Luo, C. et al. Single-cell methylomes identify neuronal subtypes and regulatory elements in mammalian cortex. Science 357, 600–604 (2017).

9. Yin, Y. et al. Impact of cytosine methylation on DNA binding specificities of human transcription factors. Science 356, eaaj2239 (2017).

10. Zhu, H., Wang, G. & Qian, J. Transcription factors as readers and effectors of DNA methylation. Nat. Rev. Genet. 17, 551–565 (2016).

11. Stricker, S. H., Koferle, A. & Beck, S. From profiles to function in epigenomics. Nat. Rev. Genet. 18, 51–66 (2017).

12. Gabel, H. W. et al. Disruption of DNA-methylation-dependent long gene repression in Rett syndrome. Nature 522, 89–93 (2015).

13. Chen, L. et al. MeCP2 binds to non-CG methylated DNA as neurons mature, influencing transcription and the timing of onset for Rett syndrome. Proc. Natl. Acad. Sci. U. S. A. 112, 5509–5514 (2015).

14. Stroud, H. et al. Early-Life Gene Expression in Neurons Modulates Lasting Epigenetic States. Cell 171, 1151–1164.e16 (2017).

15. Luo, C. et al. Single nucleus multi-omics links human cortical cell regulatory genome diversity to disease risk variants. bioRxiv 2019.12.11.873398 (2019) doi:10.1101/2019.12.11.873398.

16. Luo, C. et al. Robust single-cell DNA methylome profiling with snmC-seq2. Nat. Commun. 9, 3824 (2018).

17. Preissl, S. et al. Single-nucleus analysis of accessible chromatin in developing mouse forebrain reveals cell-type-specific transcriptional regulation. Nat. Neurosci. 21, 432–439 (2018).

18. Lee, D.-S. et al. Simultaneous profiling of 3D genome structure and DNA methylation in single human cells. Nat. Methods 16, 999–1006 (2019).

19. McInnes, L., Healy, J. & Melville, J. UMAP: Uniform Manifold Approximation and Projection for Dimension Reduction. arXiv [stat.ML] (2018).

20. Zhao, C., Deng, W. & Gage, F. H. Mechanisms and functional implications of adult neurogenesis. Cell 132, 645–660 (2008).

21. Carleton, A., Petreanu, L. T., Lansford, R., Alvarez-Buylla, A. & Lledo, P.-M. Becoming a new neuron in the adult olfactory bulb. Nat. Neurosci. 6, 507–518 (2003).

22. Tasic, B. et al. Shared and distinct transcriptomic cell types across neocortical areas. Nature 563, 72–78 (2018).

23. Yao, Z. et al. An integrated transcriptomic and epigenomic atlas of mouse primary motor cortex cell types. bioRxiv 2020.02.29.970558 (2020) doi:10.1101/2020.02.29.970558.

24. Yao, Z. et al. A taxonomy of transcriptomic cell types across the isocortex and hippocampal formation. bioRxiv 2020.03.30.015214 (2020) doi:10.1101/2020.03.30.015214.

25. Huang, Z. J. & Paul, A. The diversity of GABAergic neurons and neural communication elements. Nat. Rev. Neurosci. 20, 563–572 (2019).

26. Hodge, R. D. et al. Conserved cell types with divergent features in human versus mouse cortex. Nature (2019) doi:10.1038/s41586-019-1506-7.

27. Krienen, F. M. et al. Innovations in Primate Interneuron Repertoire. bioRxiv 709501 (2019) doi:10.1101/709501.

28. Dudek, S. M., Alexander, G. M. & Farris, S. Rediscovering area CA2: unique properties and functions. Nat. Rev. Neurosci. 17, 89–102 (2016).

29. San Antonio, A., Liban, K., Ikrar, T., Tsyganovskiy, E. & Xu, X. Distinct physiological and developmental properties of hippocampal CA2 subfield revealed by using anti-Purkinje cell protein 4 (PCP4) immunostaining. J. Comp. Neurol. 522, 1333–1354 (2014).

30. Fukaya, M., Yamazaki, M., Sakimura, K. & Watanabe, M. Spatial diversity in gene expression for VDCCgamma subunit family in developing and adult mouse brains. Neurosci. Res. 53, 376–383 (2005).

31. Phillips, H. S., Hains, J. M., Laramee, G. R., Rosenthal, A. & Winslow, J. W. Widespread expression of BDNF but not NT3 by target areas of basal forebrain cholinergic neurons. Science 250, 290–294 (1990).

32. Adamek, G. D., Shipley, M. T. & Sanders, M. S. The indusium griseum in the mouse: architecture, Timm’s histochemistry and some afferent connections. Brain Res. Bull. 12, 657–668 (1984).

33. Heimer, L. & Wilson, R. D. The subcortical projections of the allocortex: similarities in the neural connections of the hippocampus, the piriform cortex and the neocortex. Santini M, editor. Perspectives in neurobiology (1975).

34. Voorn, P., Vanderschuren, L. J. M. J., Groenewegen, H. J., Robbins, T. W. & Pennartz, C. M. A. Putting a spin on the dorsal-ventral divide of the striatum. Trends Neurosci. 27, 468–474 (2004).

35. Smith, J. B. et al. Genetic-Based Dissection Unveils the Inputs and Outputs of Striatal Patch and Matrix Compartments. Neuron 91, 1069–1084 (2016).

36. Zhang, Z. et al. Epigenomic Diversity of Cortical Projection Neurons in the Mouse Brain. bioRxiv 2020.04.01.019612 (2020) doi:10.1101/2020.04.01.019612.

37. Mukamel, E. A. & Ngai, J. Perspectives on defining cell types in the brain. Curr. Opin. Neurobiol. 56, 61–68 (2019).

38. Haghverdi, L., Lun, A. T. L., Morgan, M. D. & Marioni, J. C. Batch effects in single-cell RNA-sequencing data are corrected by matching mutual nearest neighbors. Nat. Biotechnol. 36, 421–427 (2018).

39. Hie, B., Bryson, B. & Berger, B. Efficient integration of heterogeneous single-cell transcriptomes using Scanorama. Nat. Biotechnol. (2019) doi:10.1038/s41587-019-0113-3.

40. Deneris, E. S. & Hobert, O. Maintenance of postmitotic neuronal cell identity. Nat. Neurosci. 17, 899–907 (2014).

41. Kepecs, A. & Fishell, G. Interneuron cell types are fit to function. Nature 505, 318–326 (2014).

42. Paul, A. et al. Transcriptional Architecture of Synaptic Communication Delineates GABAergic Neuron Identity. Cell 171, 522–539.e20 (2017).

43. Fornes, O. et al. JASPAR 2020: update of the open–access database of transcription factor binding profiles. Nucleic Acids Res. (2019) doi:10.1093/nar/gkz1001.

44. Cubelos, B. et al. Cux1 and Cux2 regulate dendritic branching, spine morphology, and synapses of the upper layer neurons of the cortex. Neuron 66, 523–535 (2010).

45. Oishi, K., Aramaki, M. & Nakajima, K. Mutually repressive interaction between Brn1/2 and Rorb contributes to the establishment of neocortical layer 2/3 and layer 4. Proc. Natl. Acad. Sci. U. S. A. 113, 3371–3376 (2016).

46. Arendt, D. et al. The origin and evolution of cell types. Nat. Rev. Genet. 17, 744–757 (2016).

47. Smith, J. B. et al. The relationship between the claustrum and endopiriform nucleus: A perspective towards consensus on cross-species homology. J. Comp. Neurol. 527, 476–499 (2019).

48. Crick Francis C & Koch Christof. What is the function of the claustrum? Philos. Trans. R. Soc. Lond. B Biol. Sci. 360, 1271–1279 (2005).

49. Puelles, L. Chapter 4 - Development and Evolution of the Claustrum. in The Claustrum (eds. Smythies, J. R., Edelstein, L. R. & Ramachandran, V. S.) 119–176 (Academic Press, 2014).

50. Ross, S. E., Greenberg, M. E. & Stiles, C. D. Basic helix-loop-helix factors in cortical development. Neuron 39, 13–25 (2003).

51. He, Y. et al. Improved regulatory element prediction based on tissue-specific local epigenomic signatures. Proc. Natl. Acad. Sci. U. S. A. 114, E1633–E1640 (2017).

52. Arlotta, P. et al. Neuronal subtype-specific genes that control corticospinal motor neuron development in vivo. Neuron 45, 207–221 (2005).

53. Allen, T. & Lobe, C. G. A comparison of Notch, Hes and Grg expression during murine embryonic and post-natal development. Cell. Mol. Biol. 45, 687–708 (1999).

54. Hrvatin, S. et al. PESCA: A scalable platform for the development of cell-type-specific viral drivers. bioRxiv 570895 (2019) doi:10.1101/570895.

55. Mich, J. K. et al. Epigenetic landscape and AAV targeting of human neocortical cell classes. bioRxiv 555318 (2019) doi:10.1101/555318.

56. Dimidschstein, J. et al. A viral strategy for targeting and manipulating interneurons across vertebrate species. Nat. Neurosci. 19, 1743–1749 (2016).

57. Dekker, J. & Mirny, L. The 3D Genome as Moderator of Chromosomal Communication. Cell 164, 1110–1121 (2016).

58. Ferland, R. J., Cherry, T. J., Preware, P. O., Morrisey, E. E. & Walsh, C. A. Characterization of Foxp2 and Foxp1 mRNA and protein in the developing and mature brain. J. Comp. Neurol. 460, 266–279 (2003).

59. Siddiqui, T. J. et al. An LRRTM4-HSPG complex mediates excitatory synapse development on dentate gyrus granule cells. Neuron 79, 680–695 (2013).

60. Yamawaki, N., Borges, K., Suter, B. A., Harris, K. D. & Shepherd, G. M. G. A genuine layer 4 in motor cortex with prototypical synaptic circuit connectivity. Elife 3, e05422 (2014).

61. Nieto, M. et al. Expression of Cux-1 and Cux-2 in the subventricular zone and upper layers II--IV of the cerebral cortex. J. Comp. Neurol. 479, 168–180 (2004).

62. Fanselow, M. S. & Dong, H.-W. Are the dorsal and ventral hippocampus functionally distinct structures? Neuron 65, 7–19 (2010).

63. Cembrowski, M. S., Wang, L., Sugino, K., Shields, B. C. & Spruston, N. Hipposeq: a comprehensive RNA-seq database of gene expression in hippocampal principal neurons. Elife 5, e14997 (2016).

64. Zhang, T.-Y. et al. Environmental enrichment increases transcriptional and epigenetic differentiation between mouse dorsal and ventral dentate gyrus. Nat. Commun. 9, 298 (2018).

65. Hon, G. C. et al. Epigenetic memory at embryonic enhancers identified in DNA methylation maps from adult mouse tissues. Nat. Genet. 45, 1198–1206 (2013).

66. Sansom, S. N. & Livesey, F. J. Gradients in the brain: the control of the development of form and function in the cerebral cortex. Cold Spring Harb. Perspect. Biol. 1, a002519 (2009).

67. O’Leary, D. D. M., Chou, S.-J. & Sahara, S. Area patterning of the mammalian cortex. Neuron 56, 252–269 (2007).

68. Gorkin, D. U. et al. Systematic mapping of chromatin state landscapes during mouse development. bioRxiv 166652 (2017) doi:10.1101/166652.

69. Allen Institute for Brain Science. Allen Mouse Brain Reference Atlas CCF v3. Allen Mouse Brain Reference Atlas CCF v3 http://atlas.brain-map.org (2017).

70. Schultz, M. D. et al. Human body epigenome maps reveal noncanonical DNA methylation variation. Nature 523, 212–216 (2015).

71. Amemiya, H. M., Kundaje, A. & Boyle, A. P. The ENCODE Blacklist: Identification of Problematic Regions of the Genome. Sci. Rep. 9, 9354 (2019).

72. Wolf, F. A., Angerer, P. & Theis, F. J. SCANPY: large-scale single-cell gene expression data analysis. Genome Biol. 19, 15 (2018).

73. Traag, V. A., Waltman, L. & van Eck, N. J. From Louvain to Leiden: guaranteeing well-connected communities. Sci. Rep. 9, 5233 (2019).

74. Guyon, I., Weston, J., Barnhill, S. & Vapnik, V. Gene Selection for Cancer Classification using Support Vector Machines. Mach. Learn. 46, 389–422 (2002).

75. Brodersen, K. H., Ong, C. S., Stephan, K. E. & Buhmann, J. M. The Balanced Accuracy and Its Posterior Distribution. in 2010 20th lnternational Conference on Pattern Recognition 3121-3124 (2010).

76. Lemaitre, G., Nogueira, F. & Aridas, C. K. Imbalanced-learn: A Python Toolbox to Tackle the Curse of Imbalanced Datasets in Machine Learning. J. Mach. Learn. Res. 18, 1–5 (2017).

77. Maaten, L. van der & Hinton, G. Visualizing Data using t-SNE. J. Mach. Learn. Res. 9, 2579–2605 (2008).

78. Linderman, G. C., Rachh, M., Hoskins, J. G., Steinerberger, S. & Kluger, Y. Fast interpolation-based t-SNE for improved visualization of single-cell RNA-seq data. Nat. Methods 16, 243–245 (2019).

79. Suzuki, R. & Shimodaira, H. Pvclust: an R package for assessing the uncertainty in hierarchical clustering. Bioinformatics 22, 1540–1542 (2006).

80. Grant, C. E., Bailey, T. L. & Noble, W. S. FIMO: scanning for occurrences of a given motif Bioinformatics 27, 1017–1018 (2011).

81. Wingender, E., Schoeps, T., Haubrock, M., Krull, M. & Donitz, J. TFClass: expanding the classification of human transcription factors to their mammalian orthologs. Nucleic Acids Res. 46, D343–D347 (2018).

82. Rao, S. S. P. et al. A 3D map of the human genome at kilobase resolution reveals principles of chromatin looping. Cell 159, 1665–1680 (2014).

83. Urich, M. A., Nery, J. R., Lister, R., Schmitz, R. J. & Ecker, J. R. MethylC-seq library preparation for base-resolution whole-genome bisulfite sequencing. Nat. Protoc. 10, 475–483 (2015).

84. Zhou, J. et al. Robust single-cell Hi-C clustering by convolution- and random-walk-based imputation. Proceedings of the National Academy of Sciences vol. 116 14011–14018 (2019).

85. Klopfenstein, D. V. et al. GOATOOLS: A Python library for Gene Ontology analyses. Sci. Rep. 8, 10872 (2018).

86. Zhang, H. et al. Chromatin structure dynamics during the mitosis-to-G1 phase transition. Nature 576, 158–162 (2019).

87. Srivastava, N., Hinton, G., Krizhevsky, A., Sutskever, I. & Salakhutdinov, R. Dropout: a simple way to prevent neural networks from overfitting. (2014).

88. Kingma, D. P. & Ba, J. Adam: A Method for Stochastic Optimization. arXiv [cs.LG] (2014).

89. Abadi, M. et al. Tensorflow: A system for large-scale machine learning. in 12th ${USENlX} Symposium on Operating Systems Design and lmplementation ({OSDl}$ 16) 265–283 (2016).

